# Hippocampal CA2 modulates trace fear conditioning through circuit-specific control of CA1

**DOI:** 10.64898/2026.06.05.730478

**Authors:** Thomas E. Bassett, Hui Zhang, Zorica Petrovic, Chihiro Nakamoto, Ana Cicvaric, Elizabeth M. Wood, Stephanie Rudolph, Asami Tanimura, Jelena Radulovic

**Affiliations:** Dominick P. Purpura Department of Neuroscience, Albert Einstein College of Medicine, Bronx, NY, USA; Department of Psychiatry and Behavioral Science, Albert Einstein College of Medicine, Bronx, NY, USA; Department of Biomedicine, Aarhus University, Aarhus, Denmark

## Abstract

Forming associations between temporally separated events depends on pathways linking the CA1, the subiculum (SUB), and the entorhinal cortex. The degree to which this process requires the CA2, which shares extensive connectivity with these regions, remains unknown. Using trace fear conditioning (TFC), where mice learn to associate a tone and shock separated by a temporal gap, we showed that the dCA2 contributes to TFC. Interestingly, whereas chronic dCA2 inhibition decreased cue-associated freezing, acute dCA2→dCA1 projection inhibition increased freezing. Combined with differential effects on cFos expression, this suggests that global and projection-specific perturbations of the dCA2 have distinct effects on TFC and hippocampal activity states. Fiber photometry revealed that dCA2→dCA1 activity shifted from tone responsiveness during conditioning to expected shock activation during recall, consistent with learning-associated activity remodeling. Together, these findings identify the dCA2 as a contributor to TFC and implicate dCA2→dCA1 signaling in shaping fear expression across learning and recall.

## Introduction

A central function of higher-order memory systems, shared by episodic and associative memory alike, is to preserve the temporal relationships between events separated in time.^1–3^ This ability, referred to as temporal association learning (TAL), allows memory systems to link temporally discontinuous events while preserving critical temporal details, including order, duration, and the intervals separating them.^4–6^ Collectively, TAL is essential for the transformation of discrete memory elements into meaningful memory representations, allowing past events to guide future behavior.

TAL is commonly examined in rodent models using trace fear conditioning (TFC), a behavioral paradigm in which animals learn to associate a tone with an aversive foot shock across a temporal gap known as the trace period.^4,7,8^ Previous work has implicated multiple hippocampal pathways in this process, including input from the medial entorhinal cortex (MEC) to the CA1 during TFC memory formation and the CA1 output to the SUB during memory retrieval.^9–15^ However, the contributions of other hippocampal subfields to TAL remain poorly defined. Among these, the CA2, a small hippocampal subfield situated between the CA1 and CA3, is especially well positioned to support TAL.^16–18^ The CA2 has been shown to play a critical role in bridging mnemonic delay periods during hippocampal-dependent working memory tasks while contributing to the temporal organization of activity in the CA1.^19,20^ In addition, mice with a global knockout of AVPR1B, a receptor highly enriched in CA2, exhibit impairments in temporal-order memory and object-trace-odor paired-association learning, although these findings do not definitively localize the effect to the CA2.^21^ Considered alongside the extensive connectivity of CA2 with entorhinal cortex and CA1, these observations position CA2 as a strong candidate for supporting TFC and, more broadly, TAL.^18,22,23^

Using tetanus toxin light chain (TeNT) inhibition paired with TFC in Amigo2-Cre mice (for CA2 specificity), we showed that dorsal CA2 (dCA2) inactivation impaired freezing during cue recall in TFC, while having no impact on related paradigms such as delay fear conditioning (DFC), contextual fear conditioning (CFC), or social fear conditioning (SFC).^18^ Interestingly, chemogenetic inhibition of the dCA2→dCA1 projection had the opposite effect on TFC, significantly increasing the freezing response during cue recall. cFos expression analysis of TFC animals revealed that whereas dCA2→dCA1 projection inhibition selectively disrupted CA1 cFos expression patterns, dCA2 inhibition impacted DG expression patterns, a region without a known direct connection to the CA2, suggestive of more generalized hippocampal disruption. Finally, fiber photometry recordings of the dCA2→dCA1 projection during TFC revealed an increase in activity in response to tone, as well as in the lead-up to the shock, during conditioning. During the cue test, tone-evoked activity was no longer evident, whereas activity in the expected shock window became more pronounced, consistent with a learning-associated shift in dCA2→dCA1 activity.

## Results

### dCA2 output is required for TFC

To determine whether the dCA2 contributes to TAL, we permanently blocked synaptic output from dCA2 using TeNT and looked for memory impairments in TFC (Figure 1A,B). Specifically, we bilaterally injected a Cre-dependent AAV encoding TeNT (AAV5-hSyn-DIO-TeNT-eGFP) into Amigo2-Cre mice, a transgenic mouse line with CA2-specific Cre expression, to block dCA2 synaptic transmission.^18^ A separate cohort of WT mice received the same Cre-dependent viral injection, acting as the control arm of the experiment. Three weeks following viral injection, the experimental and control cohorts were trained using TFC (see methods). Although there were no significant group differences in freezing during conditioning (Figure 1C), the global dCA2 inhibition group showed significantly reduced freezing during cue recall, particularly during the tone and trace periods (Figure 1D). In contrast, inhibition had no effect on freezing during the context test (Figure 1E). To determine whether the contribution of CA2 was specific to temporal associations rather than cued fear more generally, we used the same viral setup to train mice in DFC, a paradigm in which the tone and shock are presented concurrently without a trace period (Figure 1F,G).^24^ Notably, global dCA2 inhibition did not affect freezing during conditioning, cue recall, or the context test (Figure 1H–J), suggesting that dCA2 is engaged by the temporal demands introduced by the trace period.

**Figure 1:**
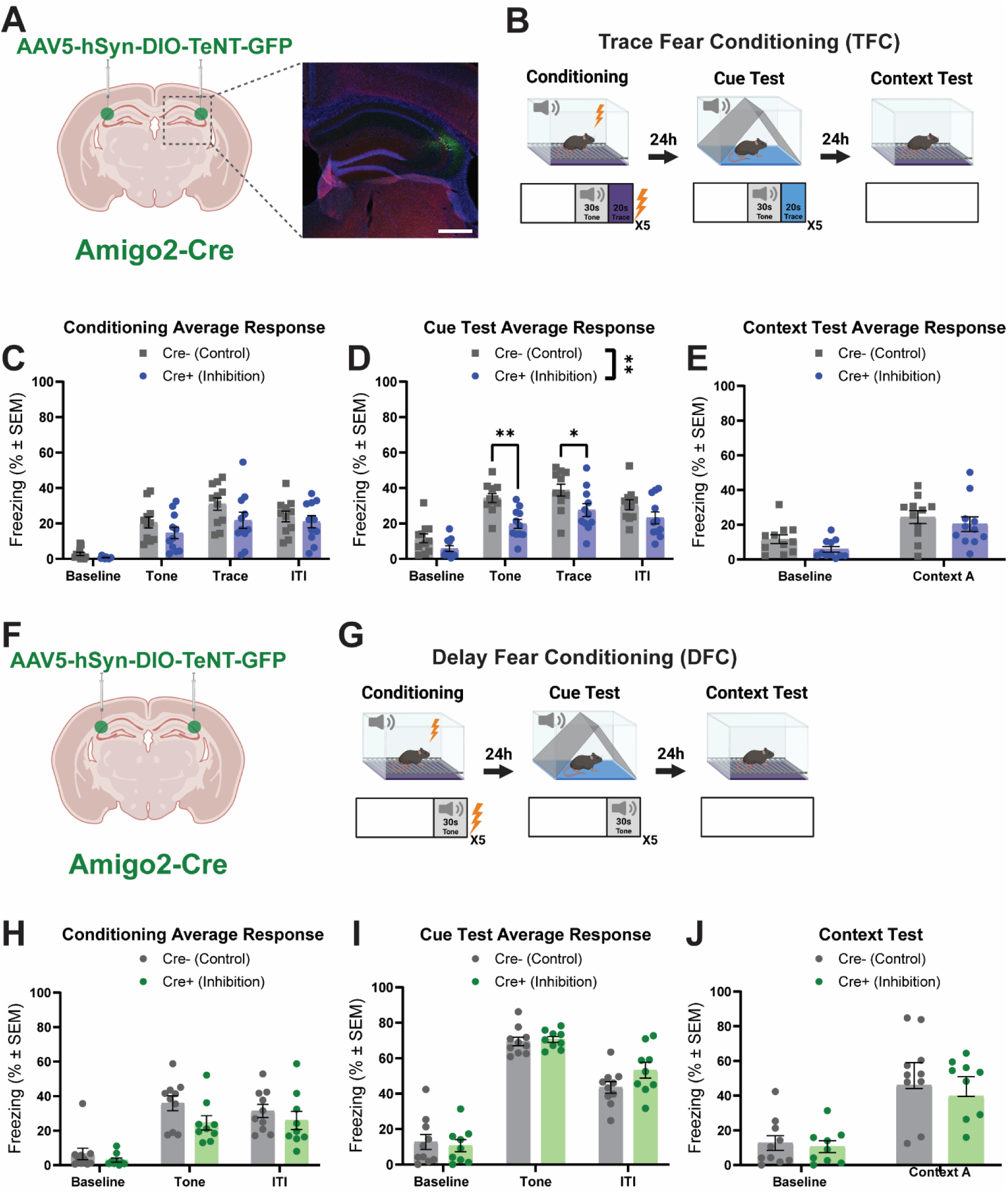
Dorsal CA2 inactivation impairs Trace Fear Conditioning. (A) Left, schematic of bilateral viral injections in Cre+ and Cre- (control) mice. Right, representative hippocampal expression in a Cre+ mouse, with hoechst in blue, virus expressing cells in green, and PCP4 (CA2 marker) in red. Scale bar – 500µm. (B) Schematic of the TFC paradigm. (C) Averaged tone, trace, and ITI freezing response calculated per animal during the conditioning trial. There was no significant difference in freezing response between the experimental and control groups (Two-way RM ANOVA; n=11; Phase×Treatment F(3,60)=1.315, P=0.2778; Phase F(3,60)=57.85, P<0.0001; Treatment F(1,20)=2.039, P=0.1687). (D) Averaged tone, trace, and ITI freezing response calculated per animal during the cue test. The experimental group froze significantly less than the control group, with multiple-comparisons finding individual significance for the baseline and tone responses (Two-way RM ANOVA; n=11; Phase×Treatment F(3,60)=2.022, P=0.1203; Phase F(3,60)=57.68, P<0.0001; Treatment F(1,20)=8.888, P=0.0074). (E) Averaged freezing response calculated per animal for the context test, with baseline freezing being the pre-tone freezing levels calculated during the cue test. There was no significant difference in freezing response between the experimental and control groups (Two-way RM ANOVA; n=11; Context×Treatment F(1,20)=0.06559, P=0.8005; Context F(1,20)=15.86, P=0.0007; Treatment F(1,20)=2.868, P=0.1059). (F) Schematic of bilateral viral injections in Cre+ and Cre- (control) mice. (G) Schematic of the delay fear conditioning paradigm. (H) Averaged tone and ITI freezing response during the conditioning trial in delay fear conditioning. There was no significant difference in freezing response between groups (Two-way ANOVA; n=10,9; Phase×Treatment F(2,34)=0.8906, P=0.4198; Phase F(2,34)=45.80, P<0.0001; Treatment F(1,17)=2.478, P=0.1339). (I) Averaged tone and ITI freezing response during the cue test trial in delay fear conditioning. There was no significant difference in freezing response between groups (Two-way ANOVA; n=10,9; Phase×Treatment F(2,34)=2.145, P=0.1327; Phase F(2,34)=203.7, P<0.0001; Treatment F(1,17)=0.7053, P=0.4127). (J) Averaged freezing response during the context test in delay fear conditioning. There was no significant difference in freezing response between groups (Two-way ANOVA; n=10,9; Context×Treatment F(1,17)=0.2120, P=0.6510; Context F(1,17)=47.49, P<0.0001; Treatment F(1,17)=0.4262, P=0.5226). For Šidák-corrected multiple comparisons: *P<0.05, **P<0.01

Given previous work implicating AVPR1B in temporal memory,^21^ we wanted to determine if AVPR1Breceptors in the dCA2 were contributing to TFC. Since AVPR1Breceptor expression is limited to the CA2 in the cornu ammonis,^25^ we tested this by infusing the AVPR1Breceptor antagonist SSR149415 into the dorsal hippocampus either before conditioning or before the cue test in TFC (Figure S2A,D).^26^ In both cases, the antagonist did not alter freezing during conditioning (Figure S2B,E) or subsequent cue retrieval (Figure S2C,F), indicating that acute AVPR1B signaling in dCA2 is not required for TFC performance.

Given the well-documented role of CA2 in social memory, particularly social recognition memory, we next asked whether this contribution extends to SFC.^17,18^ ^27,28^ To test this, we used the same dCA2 inhibition approach paired with a social fear conditioning paradigm, in which the subject mouse received an aversive foot shock upon interacting with a sex-matched conspecific in the conditioning chamber (Figure S1A,B).^29^ Interestingly, dCA2 inhibition had no impact on freezing in response to the conditioned conspecific (Figure S1C-E).

### Characterization of dCA2 inputs and outputs

Having implicated the dCA2 in TFC, we next sought to identify the circuitry that might support this contribution. However, because prior tracing studies have reported differences in the strength and topographic organization of dCA2 connectivity, particularly regarding the extent to which dCA2 engages ventral hippocampal targets^18,22,23,30–34^, we first examined the anatomical organization of its afferent and efferent projections. To examine dCA2 inputs, we unilaterally injected the dCA2 with a rabies helper AAV (AAV1-hSyn-dlox-TVA-2A-mCherry-oG), followed two weeks later by an EnvA-pseudotyped, glycoprotein-deleted rabies virus (EnvA-SAD-B19-RVΔG-emGFP-T2A-FTL), thereby restricting initial rabies uptake to dCA2 neurons. We then utilized light-sheet microscopy to generate a 3D reconstruction of the subfield’s brain-wide presynaptic inputs (Figure 2A-E). Fluorescence microscopy was utilized in combination to generate higher resolution images of select brain regions (Figure 2F-K). Light-sheet microscopy revealed that intrahippocampal inputs into the dCA2 stem primarily from the dorsal hippocampus, with much sparser soma labeling present in the intermediate hippocampus (Figure 2A-D). The dCA2 received extensive input from the dorsal CA3 (dCA3; Figure 2B,C,F), with less extensive input visible in the dorsal CA1 (dCA1), largely consistent with previous findings.^18,33,35^ Notably, we observed dense input from the dorsal SUB (dSUB; Figure 2B,C,I), a projection not previously reported in modern tracing experiments. We did not observe any DG inputs. In terms of extrahippocampal inputs, we observed extensive labelling in the MEC and lateral entorhinal cortex (LEC; Figure 2G,J), as well as in the median raphe nucleus (MR), Nucleus of the diagonal band of Broca (NDB), and in the medial septum (MS; Figure 2E,H,K).

**Figure 2:**
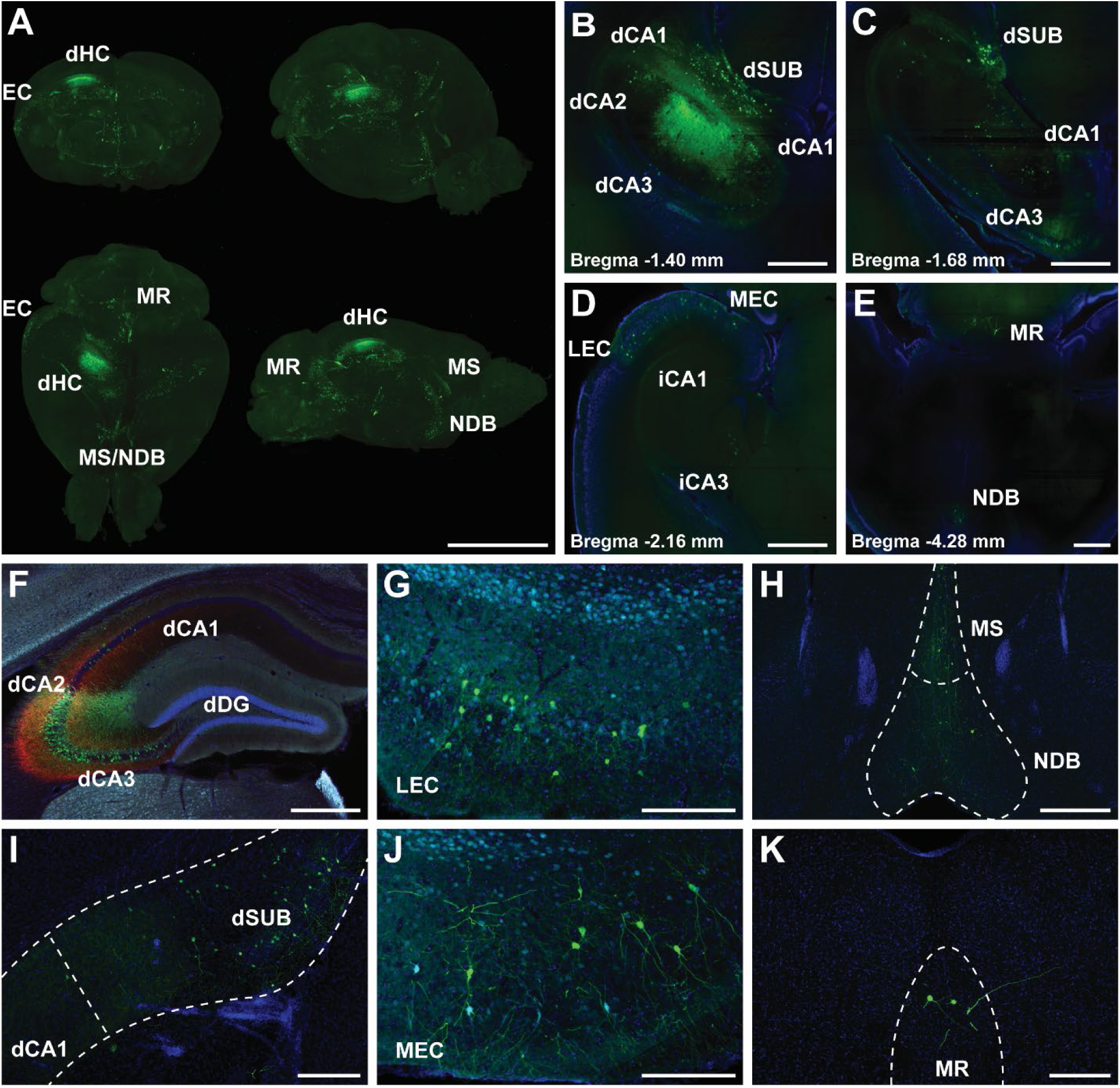
CA2 input mapping. (A) Three-dimensional reconstruction of a mouse brain used to map dorsal CA2 inputs using genetically targeted rabies tracing, with rabies labeled cells in green. (B-E) Representative horizontal sections from the 3D reconstruction, with hoechst in blue and rabies labeled cells in green. Rabies expression is depicted in the dorsal CA3, CA2, CA1, and SUB, as well as in the MEC, LEC, MR, and NDB. (F-K) Coronal fluorescence microcopy images of rabies expression, with hoechst in blue, rabies labeled cells in green, helper virus labeled cells in red, and PCP4 in cyan (n=5). (F) Dorsal hippocampus, near injection site. (G) Lateral entorhinal cortex. (H) Medial septum and nucleus of the diagonal band of broca. (I) dCA1 and dSub. (J) Medial entorhinal cortex. (K) Median raphe nucleus. Dorsal Hippocampus (dHC); Entorhinal cortex (EC); Lateral entorhinal cortex (LEC); Medial entorhinal cortex (MEC); Median raphe nucleus (MR); Nucleus of the diagonal band of Broca (NDB); Medial septum (MS) Scale Bars: A – 5mm; B-E – 1000µm; F,H – 500µm; G,I-K – 250µm

To map dCA2 outputs, we unilaterally injected a Cre-dependent synaptophysin virus (AAV1-hSyn-Flex-2A-GFP-Synaptophysin-mRuby) into Amigo2-Cre mice and used a combination of fluorescence and confocal microscopy to identify projection targets and synaptic terminal locations. Consistent with previous findings, dCA2 sent extensive bilateral projections into the dCA1, with sparser projections extending into the intermediate third of the CA1 (Figure 3A-D).^18,35^ The dCA2 also sent projections into the dCA3, with ipsilateral connections appearing denser than the contralateral ones (Figure 3A,B). Notably, we observed bilateral dCA2 projections into the dSUB, a connection that has not been consistently documented in modern CA2 tracing studies (Figure 3C,D). We did not observe labeled projections in the ventral third of the hippocampus, in contrast with previous findings.^22,28,36^

**Figure 3:**
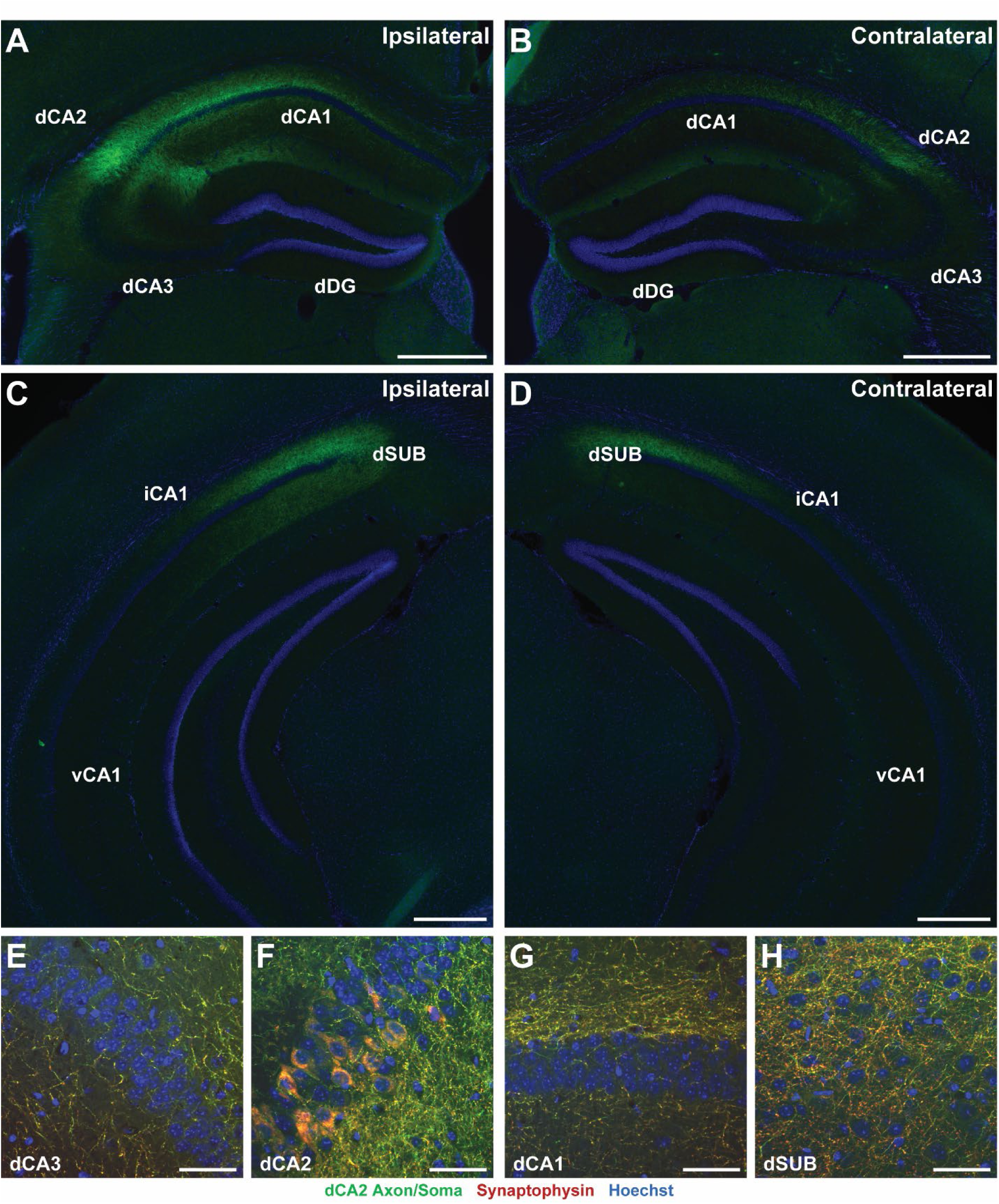
CA2 output mapping. (A) Coronal section of dorsal hippocampal viral expression. Ipsilateral hemisphere, with hoechst in blue and virus labeled cells in green. Projections stemming from the dCA2 densely innervate the dCA1, with lighter innervation in the dCA3 (n=5). (B) Coronal section of dorsal hippocampal viral expression. Contralateral hemisphere, with hoechst in blue and virus labeled cells in green. Projections into the dCA1 and dCA3, but less extensive than ipsilaterally (n=5). (C) Coronal section of ventral hippocampal viral expression. Ipsilateral hemisphere, with hoechst in blue and virus labeled cells in green. dCA2 projections heavily innervate the dSUB, and to a lesser extent the iCA1 (n=5). (D) Coronal section of ventral hippocampal viral expression. Contralateral hemisphere, with hoechst in blue and virus labeled cells in green. Projections into the dSUB and iCA1, but less extensive than ipsilaterally (n=5). (E-H) Confocal microscopy images taken in the (E) dCA3, (F) dCA2, (G) dCA1, and (H) dSUB. These images show where dCA2 projections form synapses, with hoechst in blue, virus labeled cells in green, and synaptophysin in red (n=4). Scale Bars: A-D – 500µm; E-H – 50µm

To confirm that these projections formed synapses in their target regions, we utilized confocal microscopy to visualize synaptophysin-labeled terminals in the dCA1, dCA2, dCA3, and dSUB (Figure 3E-H). Whereas labeled synapses in CA1, CA2, and CA3 largely confirmed previous findings, the presence of labeled synapses in dSUB suggests that dCA2 axons form bona fide synaptic contacts within dSUB rather than merely passing through the region.^16,22,38^

To reconcile our anterograde tracing results with prior reports of dCA2-to-ventral hippocampal connectivity, we followed up with a retrograde tracing approach. Specifically, we injected CAV2-Cre into the ventral hippocampus of Cre-dependent reporter mice (Figure S3A), which revealed a sparse population of ventral hippocampus-projecting neurons within dCA2, delineated by PCP4 (Figure S3B,C). Labeled neurons became more prominent at posterior levels of the dorsal hippocampus (Figure S3C), suggesting that this projection exists but may have been too sparse or topographically restricted to detect with our anterograde approach. Given that our viral expression was localized to the dorsal third of the hippocampus, these findings further raise the possibility that dCA2-to-ventral hippocampal projections are stronger from more intermediate CA2 levels, making injection position a potential source of variability across studies.

### Inhibiting dCA2→dCA1 projections, but not dCA2→dSUB projections, increases freezing during TFC recall

Given that the dCA1 and the dSUB are both dCA2 projection targets previously implicated in TFC, we next tested whether selective inhibition of either pathway alters freezing behavior in TFC.^9,13,15^ To selectively inhibit dCA2→dCA1 projections during conditioning, we again leveraged the Amigo2-Cre mouse line to bilaterally express an inhibitory DREADD (AAV8-hSyn-DIO-hM4Di-mCherry) exclusively in the dCA2, with cannulas implanted bilaterally right above the dCA1, such that a CNO infusion would exclusively inhibit dCA2→dCA1 projections (Figure 4A,B). WT animals that received the same viral and cannula surgeries served as the control arm of this experiment. Both experimental and control animals received a local CNO infusion through the implanted cannula 30 minutes prior to the start of behavioral testing, and GFP-tagged dextran was infused following behavioral testing to confirm appropriate cannula placement (Figure 4A). In contrast to the global inhibition experiment, dCA2→dCA1 inhibition during conditioning significantly increased freezing during cue recall, particularly during the tone, trace, and inter-trial interval (ITI) periods (Figure 4D,E). No discernible behavioral effects were observed during memory acquisition (Figure 4C). Next, we asked whether inhibiting this projection during recall would produce a similar effect. To test this, we used the same experimental setup but infused CNO during cue recall (Figure 4F,G). As during conditioning, inhibition of dCA2→dCA1 projections during the cue test increased freezing during cued recall (Figure 4H–J), suggesting that dCA2→dCA1 constrains fear expression during TFC recall.

**Figure 4:**
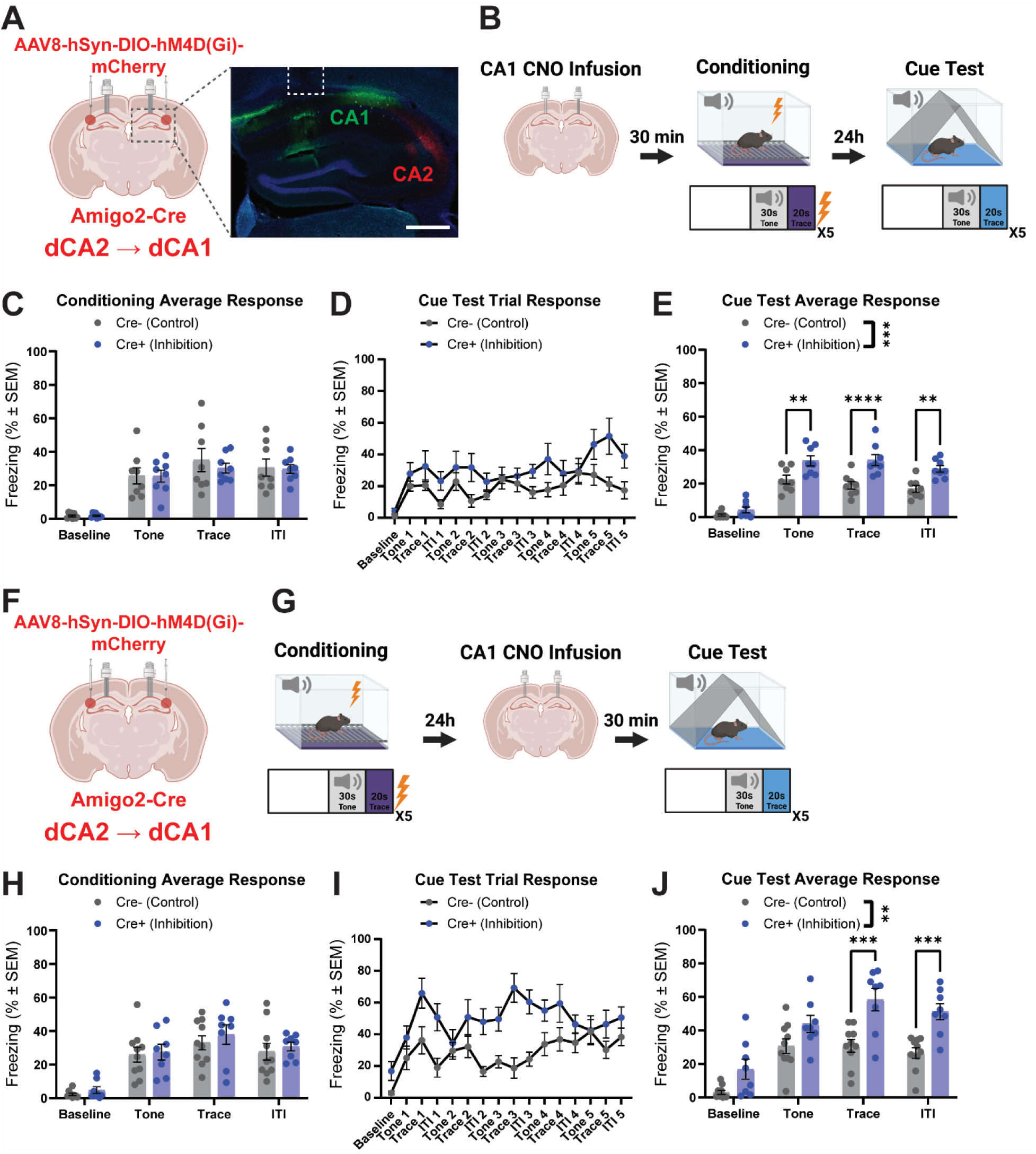
Inhibiting dCA2→dCA1 projections during conditioning and the cue test increases freezing during recall in TFC. (A) Left, schematic of bilateral viral injections in Cre+ and Cre- (control) mice. Right, representative hippocampal expression in a Cre+ mouse, with hoechst in blue, dextran conjugated to Alexa Fluor 488 in green, and virus labeled cells in red. Cannula location is denoted by the dotted white line. Scale bar – 500µm. (B) Schematic of the TFC paradigm, with CNO infusion occurring 30 minutes prior to the start of the paradigm on the conditioning day. (C) Averaged tone, trace, and ITI freezing response calculated per animal during the conditioning trial. There was no significant difference in freezing response between the experimental and control groups (Two-way ANOVA; n=8; Phase×Treatment F(3,42)=0.3781, P=0.7693; Phase F(3,42)=58.45, P<0.0001; Treatment F(1,14)=0.1019, P=0.7543). (D) Intra-trail component breakdown of freezing responses averaged across control and experimental groups during the cue test. (E) Averaged tone, trace, and ITI freezing response calculated per animal during the cue test. The experimental group froze significantly more than the control group, with multiple-comparisons finding individual significance for the tone, trace, and ITI responses (Two-way ANOVA; n=8; Phase×Treatment F(3,42)=3.752, P=0.0178; Phase F(3,42)=77.97, P<0.0001; Treatment F(1,14)=19.42, P=0.0006). (F) Schematic of bilateral viral injections in Cre+ and Cre- (control) mice. (G) Schematic of the TFC paradigm, with CNO infusion occurring 30 minutes prior to the start of the cue test. (H) Averaged tone, trace, and ITI freezing response calculated per animal during the conditioning trial. There was no significant difference in freezing response between the experimental and control groups (Two-way ANOVA; n=10,8; Phase×Treatment F(3,48)=0.1062, P=0.9561; Phase F(3,48)=43.78, P<0.0001; Treatment F(1,16)=0.4453, P=0.5141). (I) Intra-trail component breakdown of freezing responses averaged across control and experimental groups during the cue test. (J) Averaged tone, trace, and ITI freezing response calculated per animal during the cue test. The experimental group froze significantly more than the control group, with multiple-comparisons finding individual significance for the trace and ITI responses (Two-way ANOVA; n=10,8; Phase×Treatment F(3,48)=3.376, P=0.0258; Phase F(3,48)=60.36, P<0.0001; Treatment F(1,16)=14.27, P=0.0016). For Šidák-corrected multiple comparisons: **P<0.01, ***P<0.001, ****P<0.0001

To assess the relevance of dCA2→dSUB to TFC, we used the same experimental design as the dCA2→dCA1 inhibition experiment, this time implanting cannulas above the dSUB (Figure S4A,B). Inhibiting the dCA2→dSUB projection during conditioning did not significantly impact freezing during memory acquisition or during cued recall (Figure S4C-E). Similarly, dCA2→dSUB inhibition during the cue test had no impact on freezing (Figure S4F-J), suggesting that the projection is not required for the acquisition or expression of freezing behavior in TFC.

### Global dCA2 inhibition and dCA2→dCA1 projection inhibition differentially impact hippocampal cFos responses in TFC

To gain mechanistic insight into why global dCA2 inhibition and dCA2→dCA1 projection inhibition produced opposite behavioral effects in TFC, we analyzed dorsal hippocampal cFos expression following TFC under both inhibitory conditions (Figure 5A). We utilized the same targeting and inhibition strategies used during our behavioral manipulations, and trained mice only on the conditioning day of the TFC paradigm. One hour following conditioning, animals were sacrificed, brain tissue was collected, and cFos staining was performed using free-floating immunohistochemistry (see Methods). Using anatomical markers and the Allen brain atlas, we analyzed cFos expression patterns for the dCA1 and dCA3, further delineated by their distal and proximal halves, as well as in the dDG, separated into its suprapyramidal and infrapyramidal blades (Figure 5B). A standardized image processing pipeline (see Methods) was then used to quantify both the percentage of cFos-positive cells in our regions of interest, a proxy for the proportion of cells recently activated within the region, and the average intensity of cFos signal amongst positive cells, a proxy for the magnitude of activity-dependent cFos expression within recruited cells.^39,40^

**Figure 5:**
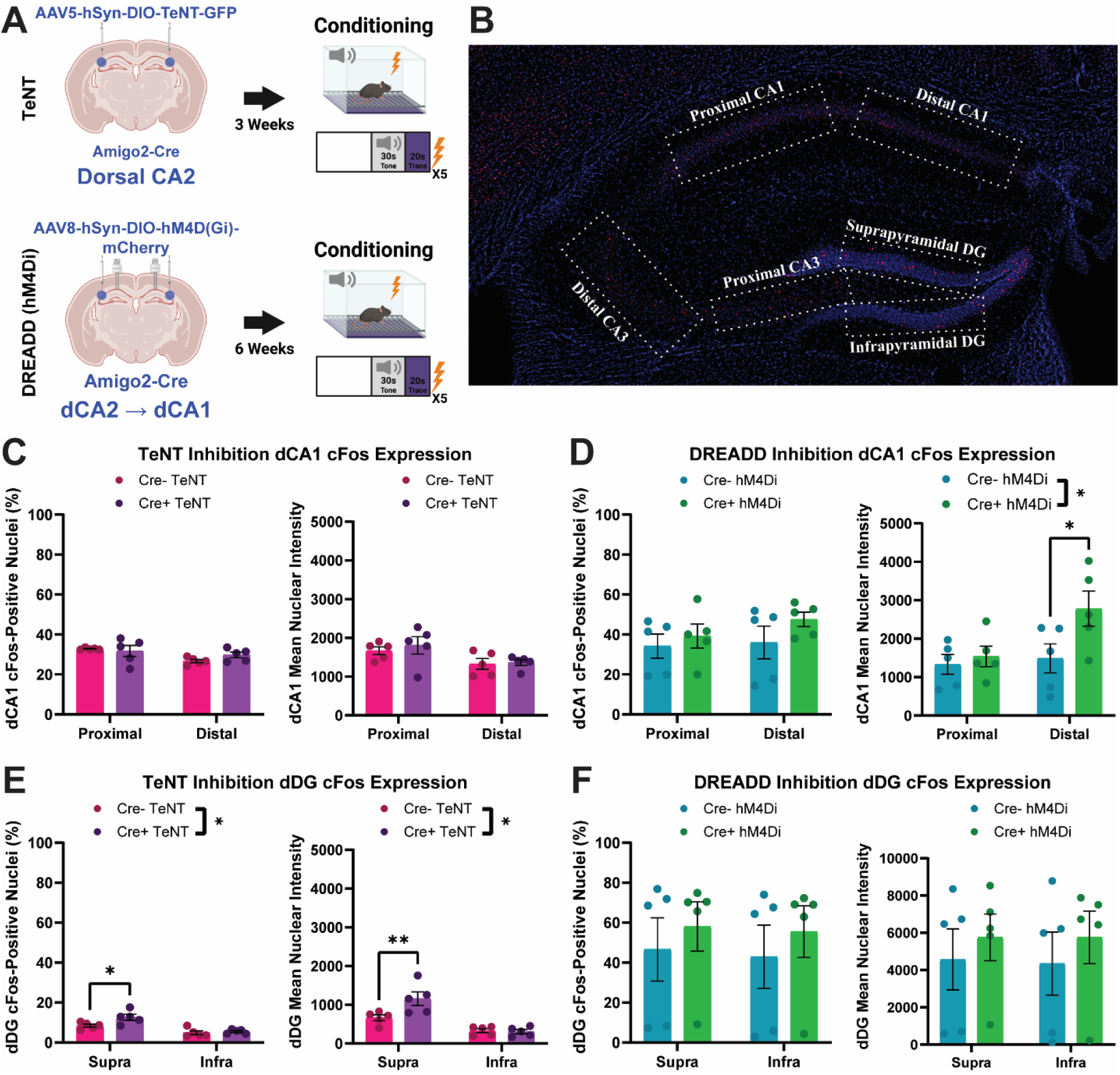
Global CA2 inhibition and dCA2→dCA1 projection inhibition differentially affect hippocampal cFos expression in TFC. (A) Experimental schematics for bilateral viral injections in Cre+ and Cre- (control) mice for global CA2 inhibition (top, TeNT) and dCA2→dCA1 projection inhibition (bottom, DREADD), paired with the conditioning day of TFC. (B) Representative microscopy image of the dorsal hippocampus showing the regions of interest used for cFos quantification. Dashed outlines indicate the analyzed hippocampal subregions, including the distal and proximal CA1, distal and proximal CA3, and suprapyramidal (Supra) and infrapyramidal (Infra) DG. (C) Left, global CA2 inhibition had no effect on the percentage of dCA1 cFos positive nuclei (Two-way ANOVA; n=5; Subregion×Treatment F(1,16)=1.629, P=0.2200; Subregion F(1,16)=6.431, P=0.0220; Treatment F(1,16)=0.3901, P=0.5410). Right, global CA2 inhibition had no effect on the mean nuclear intensity of dCA1 cFos positive nuclei (Two-way ANOVA; n=5; Subregion×Treatment F(1,16)=0.1285, P=0.7247; Subregion F(1,16)=7.250, P=0.0160; Treatment F(1,16)=0.3685, P=0.5524). (D) Left, dCA2→dCA1 projection inhibition had no effect on the percentage of dCA1 cFos positive nuclei (Two-way ANOVA; n=5; Subregion×Treatment F(1,16)=0.2825, P=0.6024; Subregion F(1,16)=0.6736, P=0.4239; Treatment F(1,16)=1.782, P=0.2006). Right, dCA2→dCA1 projection inhibition increased the mean nuclear intensity of dCA1 cFos positive nuclei relative to control (Two-way ANOVA; n=5; Subregion×Treatment F(1,16)=2.427, P=0.1388; Subregion F(1,16)=3.997, P=0.0629; Treatment F(1,16)=4.628, P=0.0471), with the effect seemingly driven by the distal portion of the subfield. (E) Left, global CA2 inhibition increased in the percentage of dDG cFos positive nuclei relative to control (Two-way ANOVA; n=5; Subregion×Treatment F(1,16)=3.214, P=0.0919; Subregion F(1,16)=29.26, P<0.0001; Treatment F(1,16)=5.828, P=0.0281). Right, global CA2 inhibition increased the mean nuclear intensity of dDG cFos positive nuclei relative to control (Two-way ANOVA; n=5; Subregion×Treatment F(1,16)=6.490, P=0.0215; Subregion F(1,16)=33.40, P<0.0001; Treatment F(1,16)=5.232, P=0.0361). In both cases, the effect was seemingly driven by the suprapyramidal blade of the subfield (F) Left, dCA2→dCA1 projection inhibition had no effect on the percentage of DG cFos positive nuclei (Two-way ANOVA; n=5; Subregion×Treatment F(1,16)=0.001604, P=0.9685; Subregion F(1,16)=0.04718, P=0.8308; Treatment F(1,16)=0.7053, P=0.4134). Right, dCA2→dCA1 projection inhibition had no effect on the mean nuclear intensity of DG cFos positive nuclei (Two-way ANOVA; n=5; Subregion×Treatment F(1,16)=0.005224, P=0.9433; Subregion F(1,16)=0.005474, P=0.9419; Treatment F(1,16)=0.7385, P=0.4028). Multiple comparisons was performed if there was a significant treatment or interaction effect For Šidák-corrected multiple comparisons: *P<0.05, **P<0.01

Within the dCA1, global inhibition during TFC did not appear to alter the number of cFos-positive cells or in the mean nuclear cFos intensity among positive cells, as compared to the control arm (Figure 5C). Notably, this was not the case for dCA2→dCA1 projection inhibition. While projection inhibition did not affect the proportion of cFos-positive nuclei within the dCA1 during TFC, there was a significant increase in the mean nuclear intensity, primarily driven by the distal segment of the subfield (Figure 5D), suggesting that projection inhibition enhanced cFos expression within recruited dCA1 neurons without increasing the size of the recruited ensemble. While not necessarily indicative of increased neuronal firing, past work examining the effects of temporary dCA2 inhibition on hippocampal activity reported an increase in sharp-wave ripples, or synchronized hippocampal populations events that prominently involve the CA1 and play a known role in TFC.^12,41^ Combined with evidence that the dCA2 exerts strong inhibitory control over the dCA1, one plausible interpretation of this cFos finding is that dCA2→dCA1 projection inhibition led to an increase in the strength or duration of activity within the already recruited population.^42^

Within the dDG, we also saw divergent cFos responses between our global and projection-specific inhibition groups. Global dCA2 inhibition resulted in an increase in the percentage of cFos-positive nuclei as well as in the mean nuclear intensity of cFos-positive cells, primarily driven by the suprapyramidal blade of the dDG (Figure 5E). Notably, this altered cFos response was not present in the projection-specific inhibition group (Figure 5F). Given the lack of a direct projection from the dCA2 to the dDG (Figure 3A,B), and the established link between chronic CA2 inhibition and hippocampal hyperexcitability,^43^ this pattern following global dCA2 inhibition may reflect a broader disruption of hippocampal network activity.

cFos analysis within the dCA3 did not reveal any changes in expression within the global dCA2 inhibition group nor within the dCA2→dCA1 projection inhibition group (Figure S5A,B). Given that the dCA3 has the sparsest body of literature linking it to TFC among the dorsal hippocampal subfields, this may provide additional evidence against its major involvement in modulating TFC freezing responses.

### Fiber photometry reveals learning-associated shifts in dCA2→dCA1 calcium dynamics during TFC

Having found that global dCA2 inhibition and selective dCA2→dCA1 projection inhibition produce distinct hippocampal cFos responses during TFC, we next sought to determine how activity in the dCA2→dCA1 pathway unfolds in real time during learning and retrieval. To accomplish this, we again leveraged the Amigo2-Cre mouse line to restrict expression of a virus encoding the calcium indicator jGCaMP7s (AAV5-hSyn-DIO-jGCaMP7s) to the dCA2 (Figure 7A). We then implanted fiber optic cannulas over the dCA1 so that we could record projection-specific dCA2→dCA1 calcium activity, a proxy for neuronal activity, during TFC acquisition and recall (Figure 7B).^44^ To better define how calcium dynamics changed as a result of learning the tone-shock association in TFC, we included a control group in which animals do not learn this association. A reduced-learning control was accomplished by extending the trace period between tone and shock to 60 seconds (TFC60), up from the 20 seconds used for our previous experiments.^45^ To validate our approach, we assessed learning by quantifying the change in freezing from baseline to the first tone presentation during the recall test. As expected, the standard TFC group learned the association (Figure 7D) whereas the TFC60 group did not (Figure S6D). Prior to analysis, raw photometry signals were processed using a standardized analysis pipeline and z-scored before being averaged across animals (Figures 7C, S6C).

To assess changes between memory acquisition and recall, we calculated the average calcium activity at tone onset, in the lead-up to shock, and in response to shock. During conditioning, both standard TFC and TFC60 animals exhibited increased dCA2→dCA1 calcium activity following tone onset across pairings (Figure 6E, 7E). However, this tone-evoked increase was maintained during the cue test only in the TFC60 group and was absent during recall in the standard TFC group (Figure 6E, 7E). We next assessed activity immediately preceding shock or expected shock by comparing sequential 10 second epochs at the end of the trace period. In the standard TFC group, dCA2→dCA1 projections exhibited increased calcium activity during the pre-shock/expected-shock period during both conditioning and the cue test, with cue-test activity diminishing across subsequent tone presentations (Figure 6F). In contrast, the TFC60 group did not exhibit an overall increase in calcium activity immediately before shock or expected shock during either conditioning or the cue test, although late-trace activity varied across pairings (Figure 7F). Finally, neither standard TFC nor TFC60 animals exhibited an overall increase in calcium activity following shock or the expected-shock period, although activity surrounding this period changed across pairings in both paradigms (Figure 6G, 7G). Together, these results suggest that dCA2→dCA1 activity is differentially organized by trace interval length: during standard TFC, activity shifts from tone onset toward the expected-shock period, whereas during TFC60, dCA2→dCA1 activity remains more strongly associated with tone onset and does not show a comparable pre-shock increase.

**Figure 6:**
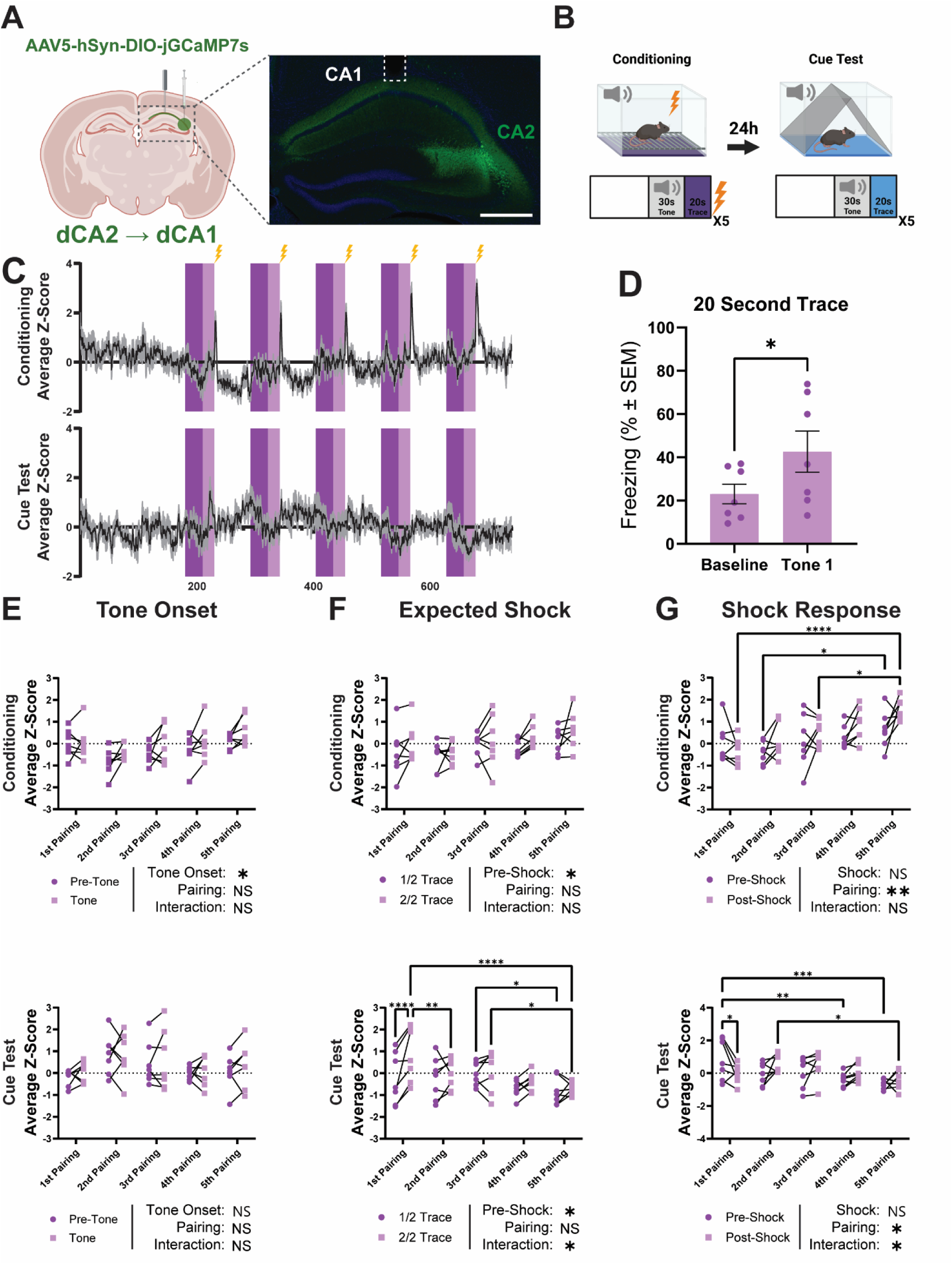
dCA2→dCA1 projections exhibit tone-evoked and pre-shock increases in calcium activity during TFC. (A) Left, schematic of viral injection and fiber placement in the dorsal hippocampus. Right, representative hippocampal expression and fiber placement image, with hoechst in blue and virus labeled cells in green. Fiber location is denoted by the dotted white line. Scale bar – 500µm. (B) Schematic of the TFC paradigm with a standard 20 second trace period. (C) Mean GCaMP traces during conditioning (top) and cue test (bottom). Dark and light purple indicate tone and trace periods. (D) Average freezing increased from baseline to the first cue-test tone, confirming tone-shock learning (Two-tailed paired t-test, P=0.0465). (E) Top, dCA2→dCA1 projections exhibited increased calcium activity following tone onset during conditioning (Two-way ANOVA; n=7; Tone×Pairing F(4,24)=0.5167, P=0.7242; Tone F(1,6)=8.151, P=0.0290; Pairing F(4,24)=2.666, P=0.0569), with the average z-score of the 10 second bin following tone onset being higher than that of the 10 second bin preceding the tone. Bottom, there was no change in calcium activity following tone onset during the cue test (Two-way ANOVA; n=7; Tone×Pairing F(4,24)=0.2033, P=0.9341; Tone F(1,6)=0.8311, P=0.3971; Pairing F(4,24)=1.403, P=0.2629). (F) dCA2→dCA1 projections exhibited increased calcium activity before expected-shock during both conditioning (top: Two-way ANOVA; n=7; Expected-Shock×Pairing F(4,24)=0.3722, P=0.8261; Expected-Shock F(1,6)=8.369, P=0.0276; Pairing F(4,24)=1.471, P=0.2422) and the cue test (bottom: Two-way ANOVA; n=7; Expected-Shock×Pairing F(4,24)=3.387, P=0.0248; Pre-Shock F(1,6)=7.440, P=0.0343; Pairing F(4,24)=2.497, P=0.0696), with the average z-score of the 10 second bin preceding the shock being higher than that of the 10 second bin starting 20 seconds before the shock. During the cue test, the significant Expected-Shock×Pairing interaction and subsequent multiple comparisons analysis suggested that this activity preceding the shock diminished across subsequent pairings. (G) dCA2→dCA1 projections did not exhibit an overall change in calcium activity following expected-shock presentation during conditioning (top: Two-way ANOVA; n=7; Shock×Pairing F(4,24)=2.296, P=0.0885; Shock F(1,6)=4.721, P=0.0728; Pairing F(4,24)=4.535, P=0.0072). However, calcium activity significantly differed across pairings, with subsequent multiple comparisons analysis suggesting increased activity during later conditioning pairings. During the cue test, dCA2→dCA1 projections also did not exhibit an overall post-shock change in calcium activity, but showed significant effects of Pairing and Shock×Pairing interaction (bottom: Two-way ANOVA; n=7; Shock×Pairing F(4,24)=3.551, P=0.0206; Shock F(1,6)=2.495, P=0.1653; Pairing F(4,24)=3.481, P=0.0223), with subsequent multiple comparisons analysis suggesting that post-shock activity changed across pairings. For Šidák-corrected multiple comparisons: *P<0.05, **P<0.01, ***P<0.001, ****P<0.0001, NS-Not Significant

**Figure 7:**
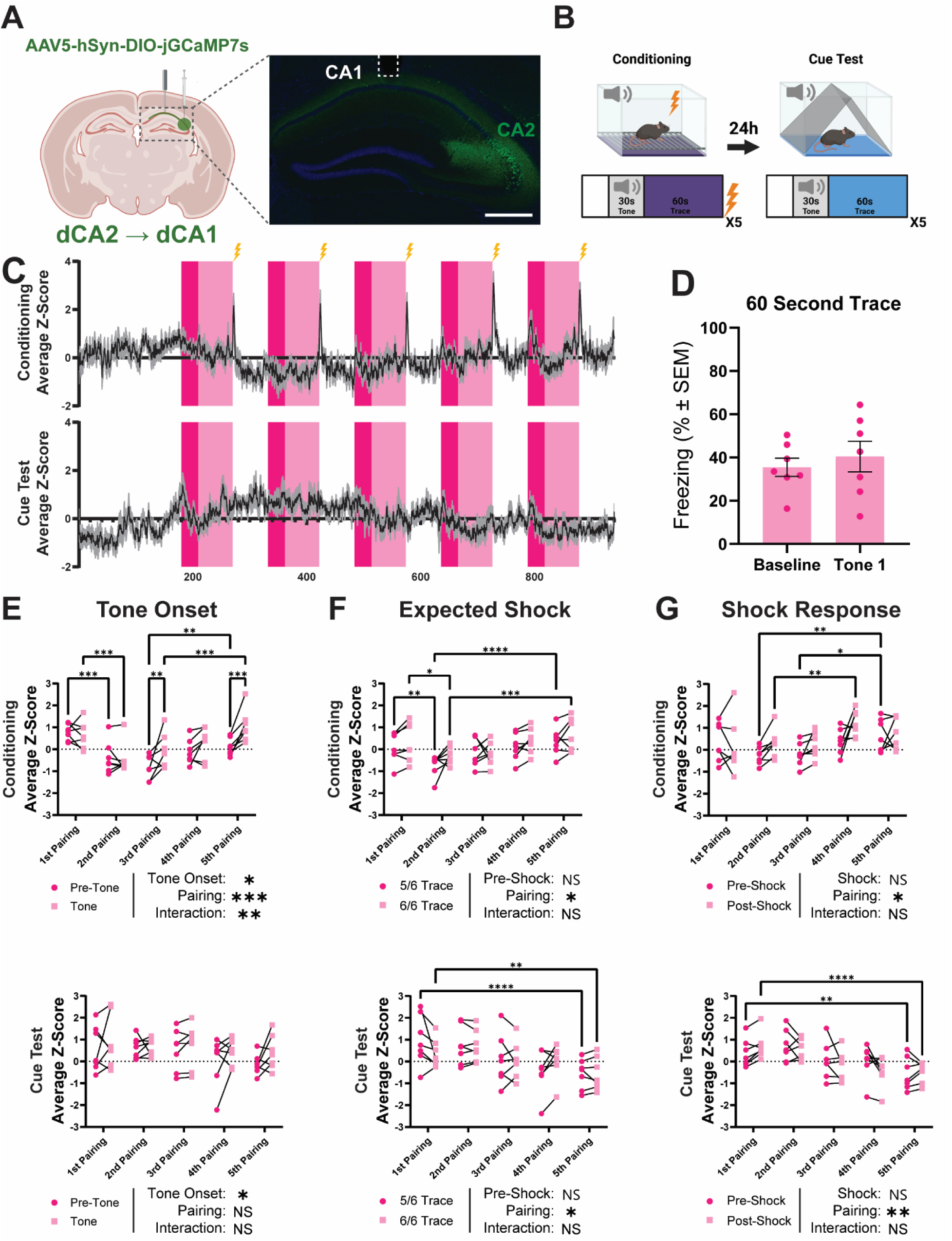
dCA2→dCA1 projections exhibit tone-evoked, but not pre-shock, increases in calcium activity during 60s-trace TFC. (A) Left, schematic of viral injection and fiber placement in the dorsal hippocampus. Right, representative hippocampal expression and fiber placement image, with hoechst in blue and virus labeled cells in green. Fiber location is denoted by the dotted white line. Scale bar – 500µm. (B) Schematic of the TFC paradigm with an extended 60 second trace period (TFC60). (C) Mean GCaMP traces during conditioning (top) and cue test (bottom). Dark and light pink indicate tone and trace periods. (D) Averaged freezing did not differ between baseline and first cue-test tone (Two-tailed paired t-test, P=0.6440). (E) Top, dCA2→dCA1 projections exhibited increased calcium activity following tone onset during conditioning, measured as a significantly higher average z-score in the 10 second bin following tone onset compared to the 10 second bin preceding the tone, with this tone-associated increase differing across pairings (Two-way ANOVA; n=7; Tone×Pairing F(4,24)=4.661, P=0.0063; Tone F(1,6)=9.542, P=0.0214; Pairing F(4,24)=8.907, P=0.0001). Subsequent multiple comparisons analysis suggested that tone-evoked activity varied across conditioning pairings. Bottom, dCA2→dCA1 projections exhibited increased calcium activity following tone onset during the cue test (Two-way ANOVA; n=7; Tone×Pairing F(4,24)=0.3691, P=0.8282; Tone F(1,6)=6.165, P=0.0476; Pairing F(4,24)=1.134, P=0.3643). (F) Top, dCA2→dCA1 projections did not exhibit an overall change in calcium activity immediately before expected-shock during conditioning, measured by comparing the average z-score of the final 10 second interval of the 60 second trace period (6/6 Trace) to that of the preceding 10 second interval (5/6 Trace), but calcium activity significantly differed across pairings (Two-way ANOVA; n=7; Expected-Shock×Pairing F(4,24)=0.9458, P=0.4548; Expected-Shock F(1,6)=3.880, P=0.0964; Pairing F(4,24)=2.869, P=0.0449). Subsequent multiple comparisons analysis suggested that late-trace activity increased across later conditioning pairings. Bottom, dCA2→dCA1 projections did not exhibit an overall change in calcium activity immediately before expected-shock during the cue test, but calcium activity significantly differed across pairings (bottom: Two-way ANOVA; n=7; Expected-Shock×Pairing F(4,24)=2.517, P=0.0679; Pre-Shock F(1,6)=0.9054, P=0.3781; Pairing F(4,24)=2.869, P=0.0449). Subsequent multiple comparisons analysis suggested that late-trace activity decreased across subsequent cue test pairings. (G) Top, dCA2→dCA1 projections did not exhibit an overall change in calcium activity following expected-shock presentation during conditioning, but calcium activity significantly differed across pairings (Two-way ANOVA; n=7; Shock×Pairing F(4,24)=2.764, P=0.0507; Shock F(1,6)=2.593, P=0.1584; Pairing F(4,24)=3.148, P=0.0325). Subsequent multiple comparisons analysis suggested increased shock-period activity during later conditioning pairings. Bottom, dCA2→dCA1 projections did not exhibit an overall change in calcium activity following the expected-shock period during the cue test, but calcium activity significantly differed across pairings (Two-way ANOVA; n=7; Shock×Pairing F(4,24)=2.137, P=0.1072; Shock F(1,6)=1.125, P=0.3296; Pairing F(4,24)=4.524, P=0.0073). Subsequent multiple comparisons analysis suggested decreased activity surrounding the expected-shock period across subsequent cue test pairings. For Šidák-corrected multiple comparisons: *P<0.05, **P<0.01, ***P<0.001, ****P<0.0001, NS-Not Significant

To determine whether freezing behavior could account for the observed fiber photometry effects, we performed correlation analyses between calcium activity and freezing behavior across all significant comparisons. While some datasets exhibited significant calcium-freezing correlations (Figure S6), such cases exhibited low explained variance (R^2^ < 0.15), indicting that freezing could only account for a small fraction of the overall signal. Thus, these findings argue against freezing behavior as the primary driver of our observed effects, although a partial contribution from movement-related activity cannot be excluded.

## Discussion

Our work has demonstrated that the CA2 contributes to TFC. Importantly, these contributions did not extend to associative memory paradigms where the conditioned and unconditioned stimuli were concurrently present, including DFC, CFC, and SFC, largely consistent with previous findings.^18,46^ This suggests that the CA2 does not broadly regulate fear conditioning, but instead supports the temporal associative demands specific to TFC. This idea is bolstered by past work that indirectly linked the CA2 to other temporal association tasks, including temporal-order memory and object-trace-odor paradigms, suggesting that the subfield’s role is not only limited to temporal associations in TFC, but rather to TAL more generally.^21,25^ We also found that pharmacological blockade of CA2 AVPR1B signaling before either conditioning or retrieval did not alter freezing in TFC, suggesting that the role of CA2 in this task is not dependent on acute vasopressin signaling under the conditions tested here.

These initial findings were particularly noteworthy because they further expand on the idea that the CA2 is more than a hub for social memory, a central focus of previous investigations.^17,18,28,31^ Recent work has implicated the subfield in multiple non-social functions, including the bridging of delay periods in working memory and the regulation of sharp-wave ripples.^19,41,47,48^ Thus, it is becoming clearer that CA2 is not accurately defined by its role in any singular behavior, but, much like its neighboring subfields, in a broader set of hippocampal computations, including those required for TAL.

One of our more unexpected findings was that our two dCA2 manipulations produced divergent effects on TFC behavior. While seemingly paradoxical at first, it is important to remember that these inhibitory manipulations were not equivalent. In the case of dCA2 inhibition with TeNT, we indiscriminately blocked all synaptic transmission stemming from dCA2 neurons, thereby disrupting the full range of downstream projections through which the subfield may contribute to TFC.^49^ Importantly, TeNT inhibition is chronic in nature, and thus the affected animals were under constant dCA2 inhibition in the days and weeks leading up to behavioral testing. Past work has shown that similar manipulations have led to hyperexcitability of the entire hippocampal network, raising the possibility that dCA2 loss is leading to network-level disruptions that render the mice unable to appropriately form temporal associations.^43^ In contrast, our DREADD-mediated dCA2→dCA1 inhibition approach was pathway-specific, targeting a projection known to engage strong feed-forward inhibitory control over CA1 activity.^16,42^ In utilizing a microinfusion system to deliver CNO directly into dCA1, rather than using systemic CNO delivery, we were able to avoid inhibiting dCA2 projections outside of dCA1 and thus minimize potential confounding effects. Additionally, DREADD inhibition is acute in nature, lasting only a few hours, minimizing the potential for longer term network effects. Taken together, these differences suggest that the opposing behavioral effects of global dCA2 inhibition and selective dCA2→dCA1 inhibition do not reflect a true contradiction but rather demonstrate that different forms of dCA2 perturbation can produce distinct effects on behavior.

Consistent with this interpretation, our cFos analyses revealed that global dCA2 inhibition and dCA2→dCA1 projection inhibition differentially affected neuronal activity in individual CA subfields. For the projection-specific manipulation, inhibition increased cFos intensity among cFos-positive cells within the distal segment of the dCA1, without affecting the overall percentage of cFos-positive cells. This could be plausibly explained by an increase in sharp-wave ripple occurrence known to result from acute CA2 inhibition loss of feed-forward inhibition from dCA2→dCA1 projections, resulting in increased ensemble activation without ensemble expansion.^41,42^ Notably, cFos was not differentially expressed in the dDG or dCA3, indicative of a more selective subfield effect. Interestingly, global dCA2 inhibition did not affect cFos expression in either the dCA1 or the dCA3, but did increase both the percentage and average intensity of cFos-positive cells in the suprapyramidal blade of the dDG. Given that the dCA2 does not directly project to the dDG, this effect is unlikely to be a direct consequence of dCA2 silencing and is instead suggestive of a broader disruption to hippocampal network activity. Because TeNT inhibition was chronic in nature, the prolonged loss of dCA2 output may have allowed compensatory adaptations to emerge within downstream hippocampal circuitry, blunting gross regional cFos differences in dCA1 and dCA3 despite persistent network dysfunction. Regardless of the mechanism, these findings support the view that global dCA2 inhibition and selective dCA2→dCA1 inhibition result in distinct hippocampal network states, helping to explain how two manipulations to the same subfield can produce divergent effects on TFC behavior.

Previous work has implicated the dCA2 in multiple hippocampal functions, notably temporal coding and the regulation of hippocampal network coordination, both of which are relevant for TAL.^19,41^ While not necessarily exclusive to this interpretation, our finding that dCA2→dCA1 inhibition increased both dCA1 cFos intensity and freezing behavior during TFC suggests that this pathway may contribute to TAL in a modulatory capacity, shaping dCA1 activity and acting to constrain the magnitude of the behavioral response during recall. This, in turn, raises the possibility that other dCA2 projections, including those targeting the dCA3 and intermediate hippocampal subfields, may make complementary contributions to TAL. Thus, it is possible that the broader contribution of dCA2 to TAL emerges not from any single output pathway alone, but from the coordinated activity of multiple downstream targets. Nonetheless, these gaps in knowledge highlight the need for future investigation into how these pathways interact to support TAL.

While largely consistent with previous reports, our anatomical findings differed in a few notable ways. For instance, we observed projections originating from the dSUB into the dCA2, which had not been reported in recent mouse tracing studies.^18,22^ This is not entirely without precedent however, as it had been reported in an older tracing study conducted in the 1980s, albeit in rabbit rather than mouse.^50^ Another notable anatomical difference was that, although previous studies have reported dCA2 projections extending into the ventral hippocampus,^28,34^ our retrograde tracing suggested that ventral hippocampus-projecting neurons represent only a sparse subset of dCA2 neurons. Consistent with a recent study^37^, these neurons also appeared biased toward the more posterior portions of the CA2, suggesting that long-axis projections may be anatomically restricted to the intermediate and ventral segments of the CA2. Differences between our findings and prior reports may reflect differences in injection location, injection volume, viral titer, or AAV serotype, especially given that CA2 connectivity varies along its anatomical axes and that CA2 pyramidal neurons show relatively high, serotype-dependent susceptibility to AAV-mediated transgene expression.^51–53^

Our fiber photometry results revealed a dynamic pattern of dCA2→dCA1 calcium activity during TFC. In the standard TFC group, which underwent a 20-s trace period and showed evidence of tone-shock learning, tone presentation increased calcium activity during conditioning but not during the cue test. On the other hand, the activity during the trace period immediately preceding the expected shock was increased during the first test and gradually decreased thereafter. By comparison, the reduced-learning TFC60 group maintained tone-evoked calcium responses during both conditioning and the cue test but did not exhibit an overall increase in activity immediately before shock or expected shock, despite pairing-dependent changes in late-trace activity. Together, these findings suggest that stronger tone-shock learning is associated with a redistribution of dCA2→dCA1 activity away from tone onset and toward the expected-shock period.

While our data cannot determine precisely what this learning-associated shift in dCA2→dCA1 activity represents, there are multiple possibilities. One possibility is that this activity reflects a modulatory signal that helps shape and constrain dCA1 responses after the association has been learned, potentially limiting maladaptive associations to irrelevant stimuli, and thus limiting maladaptive fear responses. This interpretation is supported by our finding that acute inhibition of this projection increased both freezing and distal dCA1 cFos intensity, together with previous evidence that dCA2 provides a critical source of feed-forward inhibition to dCA1.^42^ Alternatively, this activity could reflect a signal involved in bridging information across the delay period, allowing the temporally separated tone and shock to be linked. This would be consistent with prior work suggesting that CA2 contributes to TAL by helping stabilize hippocampal representations across delays, potentially through broader network interactions with other hippocampal subfields, including CA3.^19,54,55^ Regardless of the precise mechanism, these findings suggest that dCA2→dCA1 activity is not simply a static response to environmental cues, but is dynamically reorganized across learning in a manner consistent with a role in TAL.

In summary, our findings support a model in which dCA2 contributes selectively to TAL through broad hippocampal network interactions, while the dCA2→dCA1 projection acts to constrain fear expression during memory formation and recall, potentially through feed-forward inhibitory control of dCA1 activity. Beyond expanding our understanding of hippocampal circuit function, these results raise the possibility that CA2-dependent modulation of CA1 could represent a mechanism for limiting maladaptive fear responses when the underlying associations are formed across temporally discontinuous events. This could be particularly relevant to trauma-related disorders such as post-traumatic stress disorder, in which aversive experiences often unfold as temporally structured episodes where environmental cues become associated with later aversive outcomes.^56,57^ Nevertheless, future work will be needed to determine the full scope to which CA2-dependent regulation of hippocampal circuitry contributes to the encoding, generalization, or persistence of trauma-associated memories in disease-relevant models.

## Recourse availability Lead Contact

Requests for further information and resources should be directed to and will be fulfilled by the lead contact, Jelena Radulovic.

## Materials Availability

This study did not generate new unique reagents.

## Code Availability

All code associated with this work will be made publicly available via GitHub.

## Acknowledgments

We would like to thank the Cellular and Molecular Neuroimaging Core at the Albert Einstein College of Medicine for their help generating the florescence and light-sheet microscopy datasets presented in this article. We thank Vivien Prifti for her help with the behavior experiments. This work was funded by NIMH grants MH108837, MH078064, and Lundbeck Foundation grant R310-2018-3611 to J.R. Illustrations of nonscientific data were created with BioRender.com through an institutional license.

## Author Contributions

J.R., T.E.B., S.R., and A.T. designed the study; J.R. and T.E.B. wrote the manuscript; T.E.B., Z.P., and V.P. performed all behavioral experiments; T.E.B., H.Z., C.N., and A.C. performed the imaging experiments; T.E.B. and E.M.W. performed the cFos analysis; T.E.B. and A.C. performed the fiber photometry experiments; H.Z., Z.P., and E.M.W. assisted in manuscript revision.

## Supplementary Figures

**Supplemental Figure 1:**
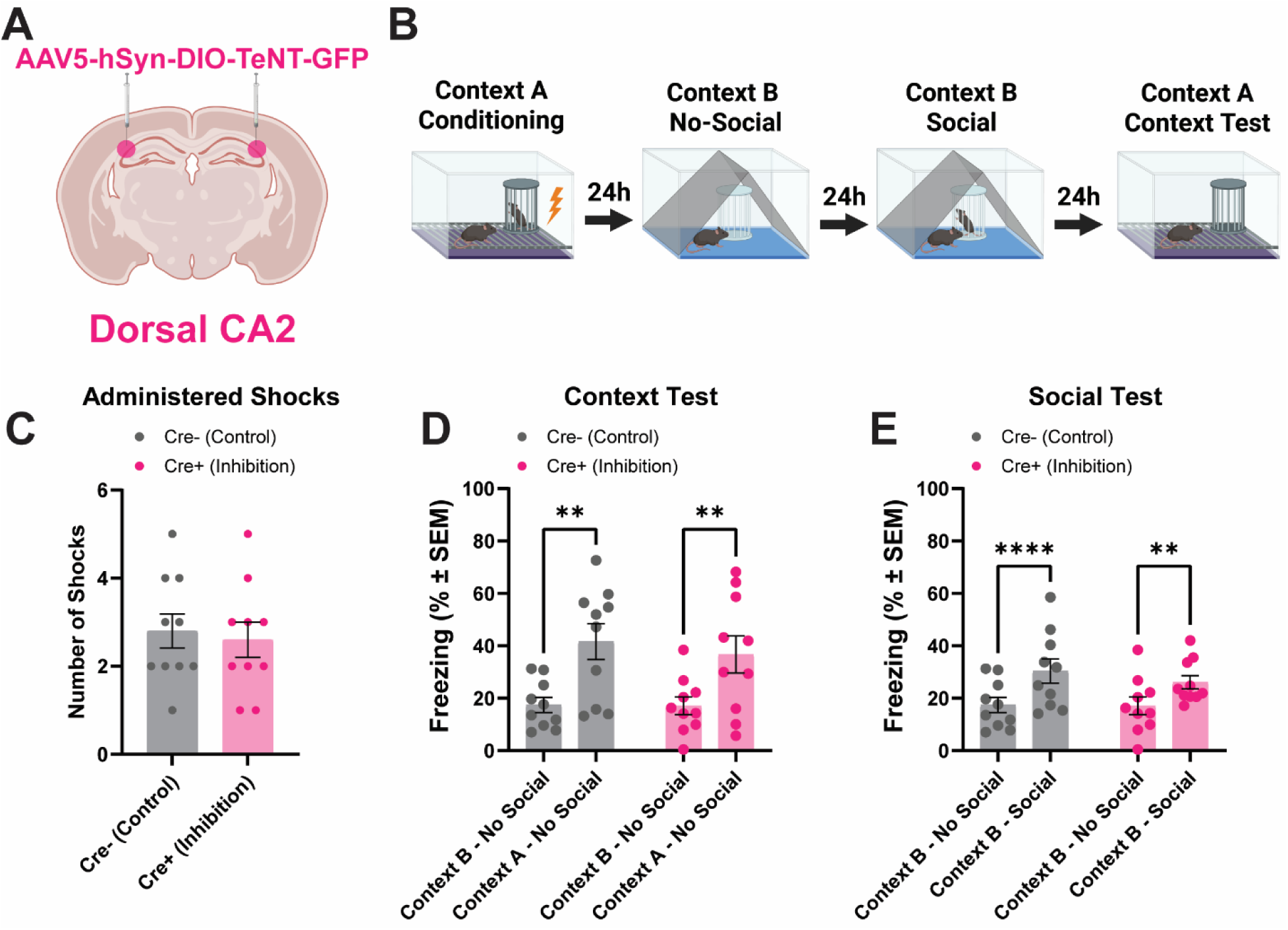
dCA2 inhibition has no effect on social fear conditioning. (A) schematic of bilateral viral injections in Cre+ and Cre- (control) mice. (B) Schematic of the social fear conditioning paradigm. (C) Number of shocks delivered to each animal, averaged across groups. There was no significant difference in the number of administered shocks between groups (n=10; Two-tailed Mann-Whitney test; P=0.7757). (D) Averaged freezing response during the context test in social fear conditioning. There was no significant difference in freezing response between groups (Two-way ANOVA; n=10; Phase×Treatment F(1,18)=0.3011, P=0.5899; Phase F(1,18)=27.96, P<0.0001; Treatment F(1,18)=0.1674, P=0.6872). (E) Averaged freezing response during the social test in social fear conditioning. There was no significant difference in freezing response between groups (Two-way ANOVA; n=10; Phase×Treatment F(1,18)=1.484, P=0.2389; Phase F(1,18)=45.56, P<0.0001; Treatment F(1,18)=0.2561, P=0.6189). For Šidák-corrected multiple comparisons: **P<0.01, **** P<0.0001

**Supplemental Figure 2:**
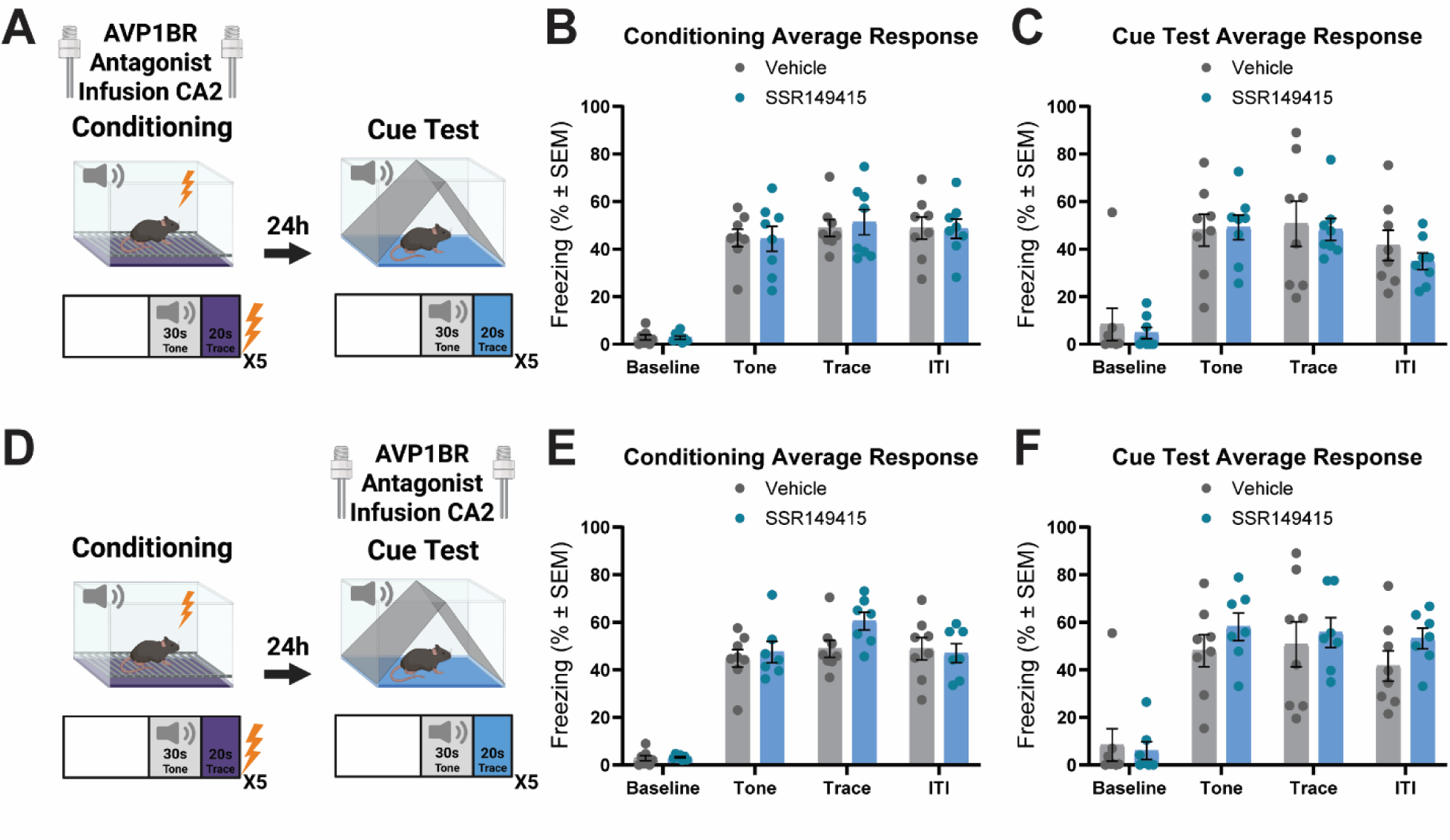
CA2 AVP1BR antagonist infusion during conditioning or retrieval did not alter freezing behavior during TFC. (A) Experimental schematic in which the AVP1BR antagonist SSR149415 or vehicle was infused into CA2 prior to the conditioning day in TFC. A shared control group was used between this experiment and the one depicted in (D). (B) Average freezing response during conditioning following pre-conditioning infusion of vehicle or SSR149415 into CA2. SSR149415 infusion did not alter freezing during baseline, tone presentation, trace period, or ITI (Two-way ANOVA; n=8; Phase×Treatment F(3,42)=0.1578, P=0.9241; Phase F(3,42)=171.0, P<0.0001; Treatment F(1,14)=0.008112, P=0.9295). (C) Average freezing response during the cue test following pre-conditioning infusion of vehicle or SSR149415 into CA2. Pre-conditioning SSR149415 infusion did not alter freezing during baseline, tone presentation, trace period, or ITI (Two-way ANOVA; n=8; Phase×Treatment F(3,42)=0.3323, P=0.8020; Phase F(3,42)=50.81, P<0.0001; Treatment F(1,14)=0.1735, P=0.6834). (D) Experimental schematic in which the AVP1BR antagonist SSR149415 or vehicle was infused into CA2 prior to the cue test in TFC. A shared control group was used between this experiment and the one depicted in (A). (E) Average freezing response during conditioning for animals subsequently infused with vehicle or SSR149415 prior to the cue test. Groups did not differ in freezing during baseline, tone presentation, trace period, or ITI (Two-way ANOVA; n=8,7; Phase×Treatment F(3,39)=2.563, P=0.0686; Phase F(3,39)=162.6, P<0.0001; Treatment F(1,13)=0.7077, P=0.4154). (F) Average freezing response during the cue test following pre-test infusion of vehicle or SSR149415 into CA2. SSR149415 infusion did not alter freezing during baseline, tone presentation, trace period, or ITI (Two-way ANOVA; n=8,7; Phase×Treatment F(3,39)=1.224, P=0.3138; Phase F(3,39)=60.09, P<0.0001; Treatment F(1,13)=0.5915, P=0.4556).

**Supplemental Figure 3:**
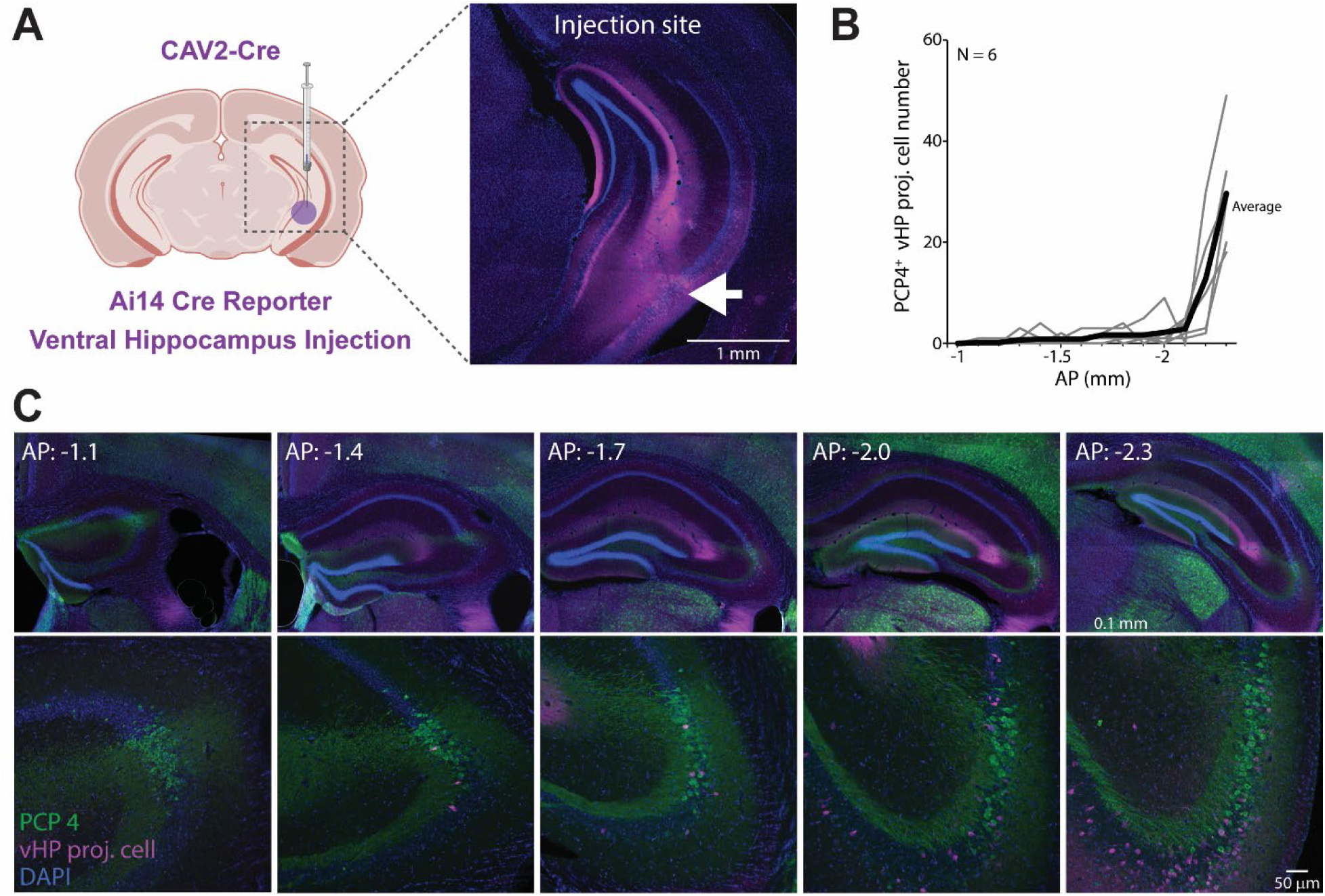
Dorsal CA2 neurons sparsely project to the ventral hippocampus. (A) Schematic of CAV2-Cre injection into the ventral hippocampus of Cre-dependent reporter mice (Ai14). Retrogradely labeled neurons projecting to the ventral hippocampus were identified by reporter expression and quantified in dorsal CA2. (B) Quantification of reporter-labeled dorsal CA2 neurons across the anterior-posterior axis of the dorsal hippocampus. Reporter-positive cells were quantified within the PCP4-positive CA2 region, revealing sparse ventral hippocampus-projecting neurons within dorsal CA2. (C) Representative dorsal hippocampal sections showing reporter-labeled ventral hippocampus-projecting neurons and PCP4 co-staining across anterior-posterior levels of dorsal CA2. DAPI is shown in blue, PCP4 in green, and reporter-labeled cells in magenta.

**Supplemental Figure 4:**
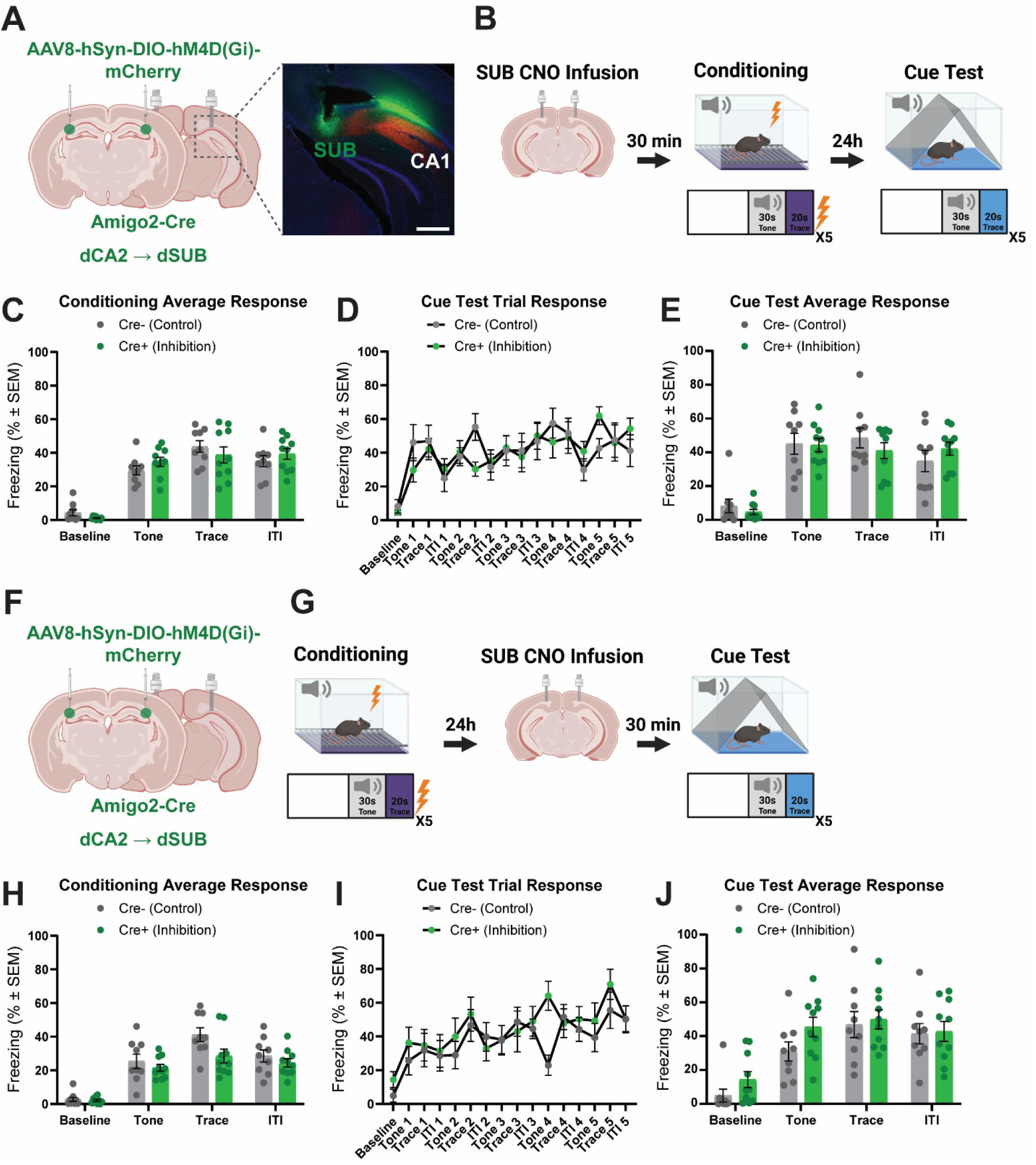
Inhibiting dCA2→dSUB projections during conditioning or cue test does not affect freezing in TFC. (A) Left, schematic of bilateral viral injections in Cre+ and Cre- (control) mice. Right, representative hippocampal expression in a Cre+ mouse, with hoechst in blue, dextran conjugated to Alexa Fluor 488 in green, and virus labeled cells in red. Scale bar – 500µm. (B) Schematic of the TFC paradigm, with CNO infusion occurring 30 minutes prior to the start of the paradigm on the conditioning day. (C) Averaged tone, trace, and ITI freezing response during the conditioning trial. There was no significant difference in freezing response between the experimental and control groups (Two-way ANOVA; n=9,10; Phase×Treatment F(3,51)=2.330, P=0.0853; Phase F(3,51)=105.7, P<0.0001; Treatment F(1,17)=0.009269, P=0.9244). (D) Intra-trail component breakdown of freezing responses averaged across control and experimental groups during the cue test. (E) Averaged tone, trace, and ITI freezing response during the cue test. There was no significant difference in freezing response between the experimental and control groups (Two-way ANOVA; n=9,10; Phase×Treatment F(3,51)=2.700, P=0.0553; Phase F(3,51)=92.62, P<0.0001; Treatment F(1,17)=0.04193, P=0.8402). (F) schematic of bilateral viral injections in Cre+ and Cre- (control) mice. (G) Schematic of the TFC paradigm, with CNO infusion occurring 30 minutes prior to the start of the cue test. (H) Averaged tone, trace, and ITI freezing response during the conditioning trial. There was no significant difference in freezing response between the experimental and control groups (Two-way ANOVA; n=9,10; Phase×Treatment F(3,51)=2.812, P=0.0485; Phase F(3,51)=80.18, P<0.0001; Treatment F(1,17)=2.475, P=0.1341). (I) Intra-trail component breakdown of freezing responses averaged across the viral control and experimental groups for the cue test. (J) Averaged tone, trace, and ITI freezing response during the cue test. There was no significant difference in freezing response between the experimental and control groups (Two-way ANOVA; n=9,10; Phase×Treatment F(3,51)=2.259, P=0.0927; Phase F(3,51)=73.01, P<0.0001; Treatment F(1,17)=0.9501, P=0.3434).

**Supplementary Figure 5:**
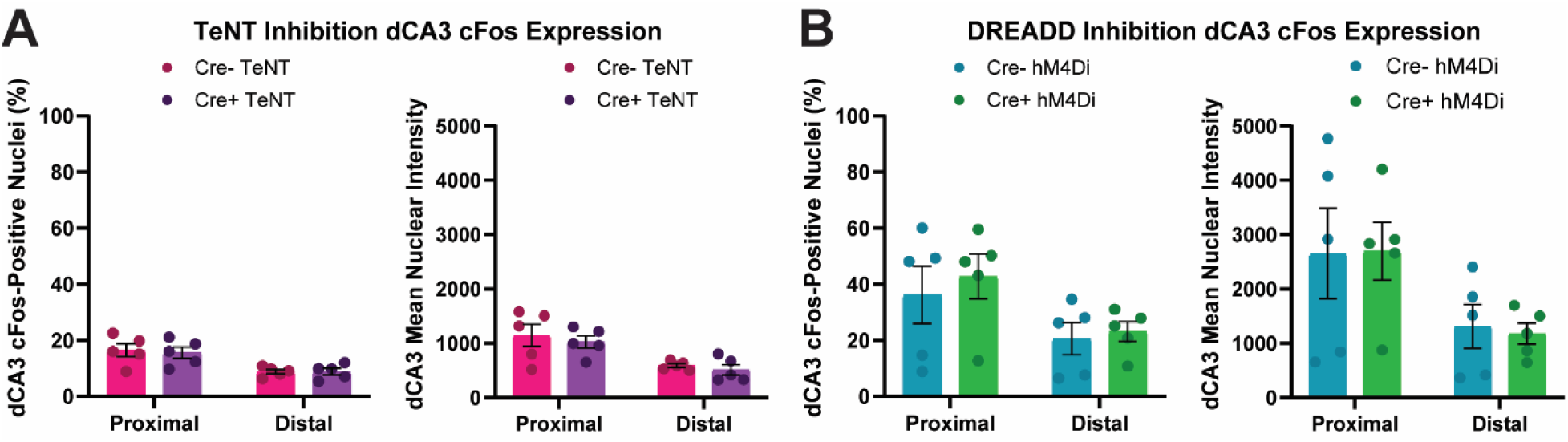
Neither global CA2 inhibition nor dCA2→dCA1 projection inhibition affect CA3 cFos expression (Related to. Figure 5). (A) Left, global CA2 inhibition had no effect on the percentage of CA3 cFos positive nuclei (Two-way ANOVA; n=5; Subregion×Treatment F(1,16)=0.06685, P=0.7993; Subregion F(1,16)=17.84, P=0.0006; Treatment F(1,16)=0.06000, P=0.8096). Right, global CA2 inhibition had no effect on the mean nuclear intensity of CA3 cFos positive nuclei (Two-way ANOVA; n=5; Subregion×Treatment F(1,16)=0.02368, P=0.8796; Subregion F(1,16)=17.20, P=0.0008; Treatment F(1,16)=0.5997, P=0.4500). (B) Left, dCA2→dCA1 projection inhibition had no effect on the percentage of CA3 cFos positive nuclei (Two-way ANOVA; n=5; Subregion×Treatment F(1,16)=0.07518, P=0.7874; Subregion F(1,16)=5.838, P=0.0280; Treatment F(1,16)=0.3848, P=0.5438). Right, dCA2→dCA1 projection inhibition had no effect on the mean nuclear intensity of CA3 cFos positive nuclei (Two-way ANOVA; n=5; Subregion×Treatment F(1,16)=0.02691, P=0.8717; Subregion F(1,16)=6.978, P=0.0178; Treatment F(1,16)=0.006961, P=0.9345).

**Supplementary Figure 6:**
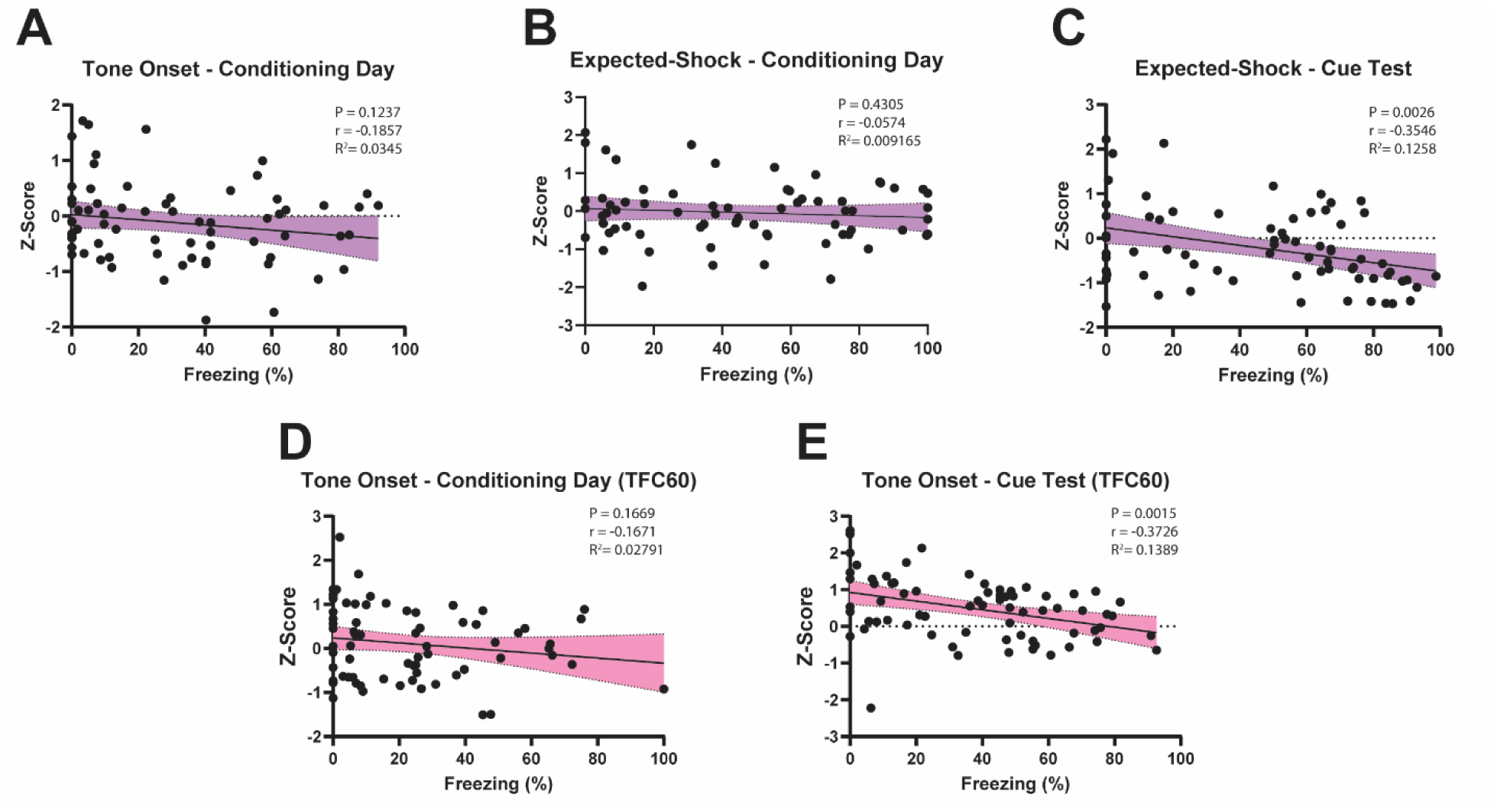
Calcium activity – freezing behavior correlation analysis. (A-E) Scatter plots showing the relationship between dCA2→dCA1 projection calcium activity and freezing behavior. P-values (P), Pearson correlation coefficients (R), and coefficient of determination (R^2^) are reported in the graphs.

## Methods

### Animal Models

We utilized male and female WT, Amigo2-Cre, and Ai14 mice throughout our experiments, all on a C57BL/6J genetic background. The Amigo2-Cre and Ai14 mouse line were obtained from the Jackson Laboratory. Amigo2-Cre mice were bred using a hemizygous breeding scheme, resulting in offspring consisting of WT and Amigo2-Cre mice, thereby allowing all experiments to be performed with littermate controls. The Amigo2-Cre mice express Cre exclusively in CA2 pyramidal neurons, thus allowing for the specific targeting of the CA2, as described previously in detail.^18^ A trio mating scheme was used, in which one hemizygous male was housed with two WT females to generate both WT and Amigo2-Cre littermates. Ai14 mice were bred using a homozygous breeding scheme, resulting in all offspring inheriting the Cre-dependent tdTomato reporter allele which limits tdTomato expression to Cre+ cells. Mice were genotyped at 5 weeks of age using primers described on the Jackson Laboratory website and used for experiments at 8 weeks of age. Litters typically consisted of 6-10 pups, with an even distribution of sex and Cre+/WT genotypes. Both male and female mice were used in behavioral, tracing, and fiber photometry experiments, with sexes evenly represented within each experiment. Mice were housed under standard conditions (12h light/dark cycle, temperature 68-72°F, humidity 30-60%) in our behavioral facilities. All animal procedures performed in this study were approved by the Albert Einstein College of Medicine’s Institutional Animal Care and Use Committee (IACUC; protocols 00001289 and 00001268) and/or the Danish national animal experiment committee (License number: 2021-15-0201-00801), in compliance with NIH standards.

### Stereotaxic surgery for viral injections and implants

Mice were anesthetized with 1.8-2.1% isoflurane and placed on a stereotaxic alignment system (Kopf model 1900). After disinfecting the surgical area, the skull was exposed, bregma was marked, and the skull was leveled to ensure accurate targeting. AAV5-hSyn-DIO-TeNT-eGFP was used for the dCA2 TeNT inhibition and cFos experiments, AAV1-hSyn-dlox-TVA-2A-mCherry-oG (helper virus) and EnvA-SAD-B19-RVΔG-emGFP-T2A-FTL (rabies virus) were used for the dCA2 input mapping experiment, AAV1-hSyn-Flex-2A-GFP-Synaptophysin-mRuby was used for the dCA2 output mapping experiment, CAV2-Cre was used for the retrograde output mapping experiment, AAV8-hSyn-DIO-hM4Di-mCherry was used for the dCA2→dCA1 inhibition, dCA2→dSUB inhibition, and cFos experiments, and AAV5-hSyn-DIO-jGCaMP7s was used for the fiber photometry experiments. With the exception of the retrograde output mapping experiment, a total of 250nl of each of the above viruses were injected into the dCA2 (AP -1.60 mm; ML ±1.80 mm; DV -1.90 mm) or the intermediate CA2 (AP -2.45 mm AP; ML ±3.00 mm, DV -2.50 mm), with inhibition experiments receiving bilateral injections, and all other experiments receiving unilateral injections. For the retrograde output mapping experiment, 60nl of virus was unilaterally injected into ventral hippocampus (AP -3.00 mm; ML +3.42 mm; DV -3.70 mm).

For behavioral experiments, viral injections were carried out using an automatic microsyringe pump controller (Micro4-WPI) and a Hamilton microsyringe (Cat # 88400). For tracing experiments, a Microliter Neuros Syringe (Cat # 65460-02) was used with the same pump controller. Viral vectors were infused over a 2 minute period, with syringes left in place for 5 minutes after the injection to allow for appropriate viral diffusion before syringe withdrawal. For the TeNT inhibition experiments, mice were given 3 weeks to allow for appropriate viral expression. For the projection-specific inhibition and fiber photometry experiments, mice were given 4 weeks to recover before cannulation surgeries were performed. For the dCA2 input mapping experiment, the helper virus was injected first and allowed to express for 2 weeks before a follow-up rabies injection was administered. The rabies virus was then allowed to express for 1 week before mice were sacrificed. For the dCA2 output mapping experiment, mice were given 4 weeks for appropriate viral expression before being sacrificed.

For the dCA2→dCA1 inhibition experiment, bilateral 26-gauge guide cannulas (Plastics One) were positioned over the dCA1 (AP -1.80 mm; ML ±1.10 mm; DV -1.5 mm) 4 weeks after viral injection surgery, and mice were given 2 week after implantation to recover before behavioral experimentation was performed. For the dCA2→dSUB inhibition experiment, the same schedule was used with two unilateral 26-gauge guide cannulas (Plastics One) positioned over the dSUB (AP -3.40 mm; ML ±1.60 mm; DV -1.7 mm), one cannula per hemisphere. For fiber photometry experiments, fiber optic cannulas (MBF Bioscience, Cat # FOC-BF-200-125, with a 200 µm core diameter and NA = 0.37) were implanted in the dCA1 (AP -1.90 mm; ML ±1.10 mm; DV -1.45 mm) 4 weeks after viral injection surgery, and mice were given 2 weeks after implantation to recover before recordings took place.

For the vasopressin antagonist experiment, two unilateral 26-gauge guide cannulas (Plastics One) positioned over the dCA2 (AP -1.80 mm; ML ±2.00 mm; DV -2.00 mm), and animals were given two weeks to recover before behavior was performed. All postoperative care was administered in accordance with our IACUC protocol.

### Trace and delay fear conditioning (TFC/DFC)

TFC was performed using an automated Med Associates fear conditioning system. On day 1 (conditioning day), mice were placed in an unfamiliar environment designated as the conditioning context (context A). This chamber was equipped with a camera to monitor movement, a wire-grid floor to deliver foot shocks, and a speaker to present auditory tones. Mice were allowed to freely explore the chamber for 180 seconds before exposure to five tone-foot-shock pairings. Each auditory tone (10 kHz, 80 dB) lasted 30 seconds and was followed by a 20-second temporal gap, referred to as the trace period. At the end of the trace period, a 2 second, 0.5 mA foot shock was delivered through the wire-grid floor. Each tone-foot shock pairing was separated by a 60 second inter-trial interval (ITI). To minimize scent-based cues, the chamber was thoroughly cleaned with 70% ethanol before each trial. On day 2 (cue test), mice were placed in a novel context (context B) that was distinct from the conditioning context. Context B differed from context A in several salient features, including floor texture, wall color, overall shape, and odor. To further differentiate the two contexts, context B was cleaned with 1% acetic acid. Together, these differences minimize generalization of contextual fear. Mice were again allowed to explore the chamber for 180 seconds before being re-exposed to the same auditory cues used during conditioning, allowing assessment of cue-associated fear memory independent of the original conditioning context. The timing of the auditory cues was consistent with conditioning day, though no shocks were administered. For certain experiments, an additional third test day was included, during which mice were returned to the conditioning context for 180 seconds to assess contextual fear memory.

Freezing behavior was recorded and quantified using the Video Freeze® software (Med Associates Inc., Fairfax, VT). Animals were selected at random to have their automated freezing scores validated through manual scoring, conducted by experimenters blinded to both treatment groups and experimental design. All behavioral experiments were conducted between 10:00 am and 5:00 pm. DFC was conducted using the same equipment and protocols as previously described for TFC. The only notable difference was that the shock was administered during the last 2 seconds of the tone, eliminating the trace period.

### Social fear conditioning (SFC)

SFC was conducted in a TSE Systems fear-conditioning chamber equipped with a wire-grid floor for foot-shock delivery and a camera for behavioral recording. On day 1, subject mice were placed in the conditioning chamber and allowed to habituate for 180 s. An empty conspecific cup was positioned in one corner of the chamber, where the social stimulus mouse would later be placed. To minimize odor cues, chambers were cleaned with 70% ethanol between animals. After the initial 180 s habituation period, the subject mouse was briefly removed, a sex-matched conspecific was placed inside the conspecific cup, and the subject mouse was returned to the chamber. The conspecific cup allowed visual, olfactory, auditory, and limited tactile interaction while preventing direct fighting and protecting the stimulus mouse from foot shock. Reintroduction of the subject mouse initiated a second 180 s trial, during which the subject received a 2 s, 0.7 mA foot shock each time it socially interacted with the stimulus mouse. Social interaction was defined as physical contact, with an experimenter blinded to treatment groups and experimental design initiating each shock. The total number of shocks delivered was recorded for each animal and capped at five. The same stimulus mouse was used throughout the paradigm for a given subject mouse. Subsequent testing days were separated by 24 hours.

On day 2, conditioned subject mice were placed in a novel context, differentiated as described for the TFC protocol. A different conspecific cup from that used on day 1 was present in the chamber but remained empty throughout the 180 s trial. On day 3, the subject mouse was returned to the day 2 context for a 180 s re-exposure trial, during which the same stimulus mouse from day 1 was placed in the conspecific cup. The difference in freezing between day 2 (no social stimulus) and day 3 (social stimulus present) was used to assess whether an association had formed between foot shock and the presence of the conspecific. On day 4, mice were returned to the original conditioning context for a final 180 s trial performed in the absence of the social stimulus mouse, allowing assessment of contextual fear memory. Freezing behavior was quantified using ezTrack.^58^

### Immunohistochemistry and microscopy

For fluorescence and confocal microscopy experiments, mice were anesthetized by intraperitoneal injection of Avertin (240 mg/kg) and transcardially perfused with 150 mL of ice-cold 4% paraformaldehyde in phosphate buffer (pH 7.4). Brains were then removed, post-fixed in the same fixative for 48 h, and cryoprotected by sequential immersion in 10%, 20%, and 30% sucrose in phosphate buffer for 24 h each. The brains were frozen, and 40 μm sections were prepared for free-floating immunohistochemistry using a cryostat (Leica CM3050 S), as previously described.^59^ Primary antibodies against mCherry (1:1000; Abcam ab167453) and GFP (1:2000; Abcam ab13970) were used to visualize viral expression; PCP4 (1:500; SYSY 480004) was used to delineate CA2; and cFos (1:3000; SYSY 226308) was used to detect neuronal cFos expression. Primary antibodies were used in conjunction with secondary antibodies from Jackson ImmunoResearch (1:500 each): Alexa Fluor 488 anti-chicken (Cat. #703-545-155), Alexa Fluor 594 anti-rabbit (Cat. #711-585-152), and Alexa Fluor 594 anti-guinea pig (Cat. #706-585-148). Hoechst (1:3000; Thermo Fisher Scientific H3570) was used as a nuclear counterstain. Sections were mounted and coverslipped using Vectashield (Vector Laboratories Cat. #H-1700). For fluorescence microscopy experiments, sections were imaged using a Zeiss Axio Scan.Z1 microscope at 10x magnification. For confocal microscopy experiments, sections were imaged using an Olympus Fluoview FV10i microscope at 60x magnification.

### 3D whole-brain light-sheet microscopy

For whole-brain light-sheet microscopy experiments, mice were anesthetized by intraperitoneal injection of Avertin (240 mg/kg) and transcardially perfused with ice-cold 4% paraformaldehyde in phosphate buffer. Brains were removed, post-fixed, and processed using the LifeCanvas Technologies SHIELD and SmartBatch+ pipeline (Lifecanvas Technologies Cat#SH-250). Briefly, samples were incubated in SHIELD-OFF solution, followed by SHIELD-ON crosslinking at 37°C for 24 hours. Samples were then delipidated using LifeCanvas delipidation buffer and a SmartBatch+ device (Lifecanvas Technologies Cat#DB). Prior to active delipidation, brains were incubated in delipidation buffer for 24 hours at 45°C with light shaking. Active delipidation was performed in a SmartBatch+ clearing cup containing delipidation buffer, with conduction buffer in the device chamber (Lifecanvas Technologies Cat#CB), using clearing mode at 42°C, 40 V, and a 1250 mA current limit for approximately 30 hours.

Cleared brains were immunolabeled using the SmartBatch+ electrophoretic labeling system. Samples were pre-incubated in SmartBatch+ Primary Sample Buffer for 2 days, refreshed in primary sample buffer for at least 3 hours, and transferred to staining cups containing primary sample buffer and primary antibodies. Primary antibodies against mCherry (40 µg; Abcam ab167453) and GFP (40 µg; Abcam ab13970) were used. Primary labeling was performed at 30°C, 90 V, and a 350 mA current limit for 18 hours. Labeling was continued as needed until the sample cup solution reached pH ≤ 8.3. Samples were then washed in PBSN at room temperature with light shaking and fixed overnight in 4% PFA. Secondary labeling was performed using Alexa Fluor 488 anti-chicken (80 µg; Jackson ImmunoResearch Cat. #703-545-155) and Alexa Fluor 594 anti-rabbit (80 µg; Jackson ImmunoResearch Cat. #711-585-152). Samples were incubated in secondary sample buffer at 37°C for 6–8 hours, transferred to staining cups containing secondary sample buffer, secondary antibodies, and SYTOX Deep Red nuclear dye (50 µL; Thermo Fisher Scientific, S11381), and labeled at 30°C, 90 V, and a 400 mA current limit for 12 hours. Samples then underwent electrophoretic washing in secondary sample buffer for 6 hours, followed by PBSN washes, overnight fixation in 4% PFA, and a final PBSN wash for at least 6 hours. Fluorescent samples were protected from light throughout processing.

Following immunolabeling, samples were refractive index matched using EasyIndex (RI = 1.52; LifeCanvas Technologies Cat#EI-100). Samples were incubated in 50% EasyIndex/50% distilled water at 37°C, followed by 100% EasyIndex at 37°C until transparent. Samples were mounted in 2% ultra-low melting point agarose prepared in EasyIndex and equilibrated in EasyIndex prior to imaging. Whole-brain image volumes were acquired using a SmartSPIM axially swept light-sheet microscope (LifeCanvas Technologies).

### Quantification of immunohistochemistry images

cFos expression was quantified from fluorescence microscopy images. Immunohistochemistry staining was performed in batch, and all images were acquired using identical microscopy settings. Sections were selected based on their posterior distance from bregma, with sections approximately 1.8 mm posterior to bregma included in the analysis. Sections were then compared against the Allen Brain Atlas to identify the primary cell layer of the dCA1, dCA3, and dDG. The dCA1 and dCA3 were further divided into their distal and proximal halves, whereas the dDG was divided into its suprapyramidal and infrapyramidal blades. These regions of interest were then cropped, with cropping dimensions being consistent across sections. The cropped images were then processed using a custom Fiji (ImageJ) macro.^60^ Briefly, nuclei were segmented from the Hoechst channel following background correction, contrast enhancement, smoothing, thresholding, and watershed separation. Nuclear ROIs were identified by particle analysis and applied to the cFos channel for intensity measurement. Local background was estimated from a perinuclear ring surrounding each nucleus, and background-subtracted cFos intensity was calculated for each ROI. Nuclei exceeding a fixed background-subtracted intensity threshold were classified as cFos-positive. Statistics including the percentage of cFos-positive nuclei and the mean intensity of cFos-positive nuclei were then generated for each image.

For the retrograde output mapping experiment, the dCA2 was delineated using a PCP4 antibody, and thus ventral hippocampus projecting dCA2 neurons were identified through the colocalization of tdTomato and PCP4. Double positive neurons in the hippocampus were manually counted using Fiji (ImageJ).^60^

### Cannula infusions

For the dCA2→dCA1 and dCA2→dSUB inhibition experiments, 1 mM water-soluble CNO (Sigma-Aldrich SML2304) was administered to animals 30 min before behavioral testing, as described previously.^28^ Briefly, animals were placed under light anesthesia (2% isoflurane), and the dummy cap sealing the cannula was removed. An injector was then inserted into the cannula and CNO was infused over a period of 30 seconds using a syringe pump (Chemyx Inc.; Fusion 400), with 500 nl per hemisphere infused in the dCA1 experiment and 300 nl per hemisphere infused in the dSUB experiment. Following infusion, the injector was left in place for 1 minute to allow the CNO to diffuse. Following removal of the injector, the dummy cap was replaced, and the animal was allowed to recover prior to testing. CNO was administered to both experimental and control groups in these experiments. To confirm accurate cannula placement and local tissue targeting, dextran conjugated to Alexa Fluor 488 (ThermoFisher D22910) was infused through the cannula 30 minutes prior to transcardial perfusion.

For the vasopressin antagonist experiment, the same general protocol used for the CNO experiments were used here. SSR149415 was used as our avp1br antagonist, which we injected bilaterally at a volume of 250nl per hemisphere at a concentration of 5 µM, in accordance with previous studies.^26^

### Fiber Photometry

Neuronal calcium dynamics were monitored in freely moving animals by fiber photometry, as described previously.^61^ Briefly, recordings were obtained with a Neurophotometrics FP3002 instrument using alternating 470 nm and 415 nm excitation, corresponding to the GCaMP-dependent and isosbestic control channels, respectively.^62,63^ Illumination was delivered through a 200 μm optical fiber with a numerical aperture of 0.37 (FOC-BF-200-125, Neurophotometrics), and output power at the fiber tip was kept stable across recording sessions within a range of 10-30 μW. Fluorescence emission was transmitted back through the same fiber, then filtered and directed onto a BlackFly CMOS camera for detection. Signals were collected in an interleaved manner at 60 frames/s. Behavioral synchronization was achieved through a TTL signal sent simultaneously from the fear-conditioning apparatus (Med Associates Inc.). Bonsai was used to acquire and record data across systems.^64^ To limit autofluorescence originating from the patch cord, the 470 nm channel was turned on for at least 15 hours before data collection. Before behavioral experiments, mice were handled and acclimated to tethering in the home cage for 3 consecutive days.

Fiber photometry signals were analyzed using GuPPy, a Python-based toolbox developed for this purpose.^65^ The 415 nm isosbestic channel was used as a reference to account for calcium-independent fluctuations, including motion-related artifacts, autofluorescence, and signal decay due to photobleaching, as it exhibits bleaching kinetics comparable to those of the calcium-dependent channel. Relative fluorescence changes were calculated as ΔF/F according to the formula: (F_observed − F_fitted) / F_fitted.^66^ The resulting traces were then high-pass filtered and converted to z scores using the session-wide mean and standard deviation, as follows: 𝑧 = (Δ𝐹/𝐹 − 𝜇_Δ_𝐹/𝐹)/𝜎_Δ_𝐹/𝐹. This normalization approach enabled comparison and pooling of recordings obtained from different animals and across testing sessions. TTL timestamps were used to align calcium recordings with the different phases of TFC. Activity periods corresponding to key timepoints in TFC were binned in 10 second intervals and compared to identify changes in overall activity.

### Quantification and Statistical Analysis

Statistical analyses were performed using GraphPad Prism. In inhibition experiments, freezing behavior was analyzed using a two-way repeated-measures ANOVA, with Treatment (Cre+ vs. WT) and TFC phase (baseline, tone, trace, and ITI) as factors. Šidák correction was used for multiple comparisons analysis. For the cFos experiments, the percentage of positive cells and mean fluorescence intensity were analyzed by two-way ANOVA with Treatment (Cre+ vs. WT) and Subregion (distal vs. proximal or suprapyramidal vs. infrapyramidal) as factors. Multiple-comparisons analyses were performed only when a significant treatment or interaction effect was detected, and between-treatment comparisons within subregions were conducted using Šidák correction. For fiber photometry experiments, activity changes were analyzed using a two-way repeated-measures ANOVA, using the TFC phase (tone, pre-shock, and shock) and pairing as factors. Šidák correction was used for multiple comparisons analysis.

Calcium activity – freezing behavior correlation analysis was conducted using Pearson’s correlation analysis. For calcium activity comparisons in which there was a statistically significant difference, the mean z-scored calcium activity from that analysis was paired and correlated with freezing percentages from those same epochs. This was done for each pairing and for each animal. r, R^2^, and P values were reported for each analysis.

